# Decoding the Functional Interactome of Non-Model Organisms with PHILHARMONIC

**DOI:** 10.1101/2024.10.25.620267

**Authors:** Samuel Sledzieski, Faith Ocitti, Charlotte Versavel, Rohit Singh, Kapil Devkota, Lokender Kumar, Polina Shpilker, Liza Roger, Jinkyu Yang, Nastassja Lewinski, Hollie Putnam, Judith Klein-Seetharaman, Bonnie Berger, Lenore Cowen

## Abstract

Despite the widespread availability of genome sequencing pipelines, many genes remain part of the genome’s “dark matter,” where existing inference tools cannot even begin to guess the biological function of their proteins from sequence alone. This challenge is especially pronounced in organisms that are highly evolutionarily distant from well-studied models, where homology-based methods break down. Here, we describe PHILHARMONIC, a computational method that combines deep learning–based *de novo* protein interaction network inference with robust unsupervised spectral clustering and remote homology to illuminate functional organization in any non-model organism. From only a sequenced proteome, we show PHILHARMONIC generates meaningful functional hypothesis for protein functions, functional communities, and higher-order network structure. We validate its performance using experimental gene expression and pathway data in *D. melanogaster*, and we demonstrate its broad utility by analyzing temperature sensing and stress response pathways in the reef-building coral *P. damicornis* and its algal symbiont *C. goreaui*. PHILHARMONIC provides a general-purpose engine for functional discovery and biological hypothesis generation in non-model organisms, enabling systems-level insights across the full diversity of life.

---

Protein-protein interaction networks are a fundamental resource for modeling molecular and cellular function, and a large and sophisticated toolbox has been developed to leverage their structure and topological organization to predict the functional roles of under-studied genes, proteins, and pathways. Common functional inference workflows leverage known protein-protein interaction (PPI) networks to cluster genes and proteins into functionally-enriched modules [1–3], or use network-propagation to directly infer protein labels based on the graph-theoretic structure [4, 5]. However, the overwhelming majority of experimentally-determined physical protein interactions from which such networks are constructed come from a small number of well-studied model organisms (Figure 1d, Appendix A.1). Indeed, most species lack even a single experimentally-determined interaction in databases such as BioGRID [6] and STRING [7], much less a network complete enough to enable the analysis of cellular function.

**Figure 1:**
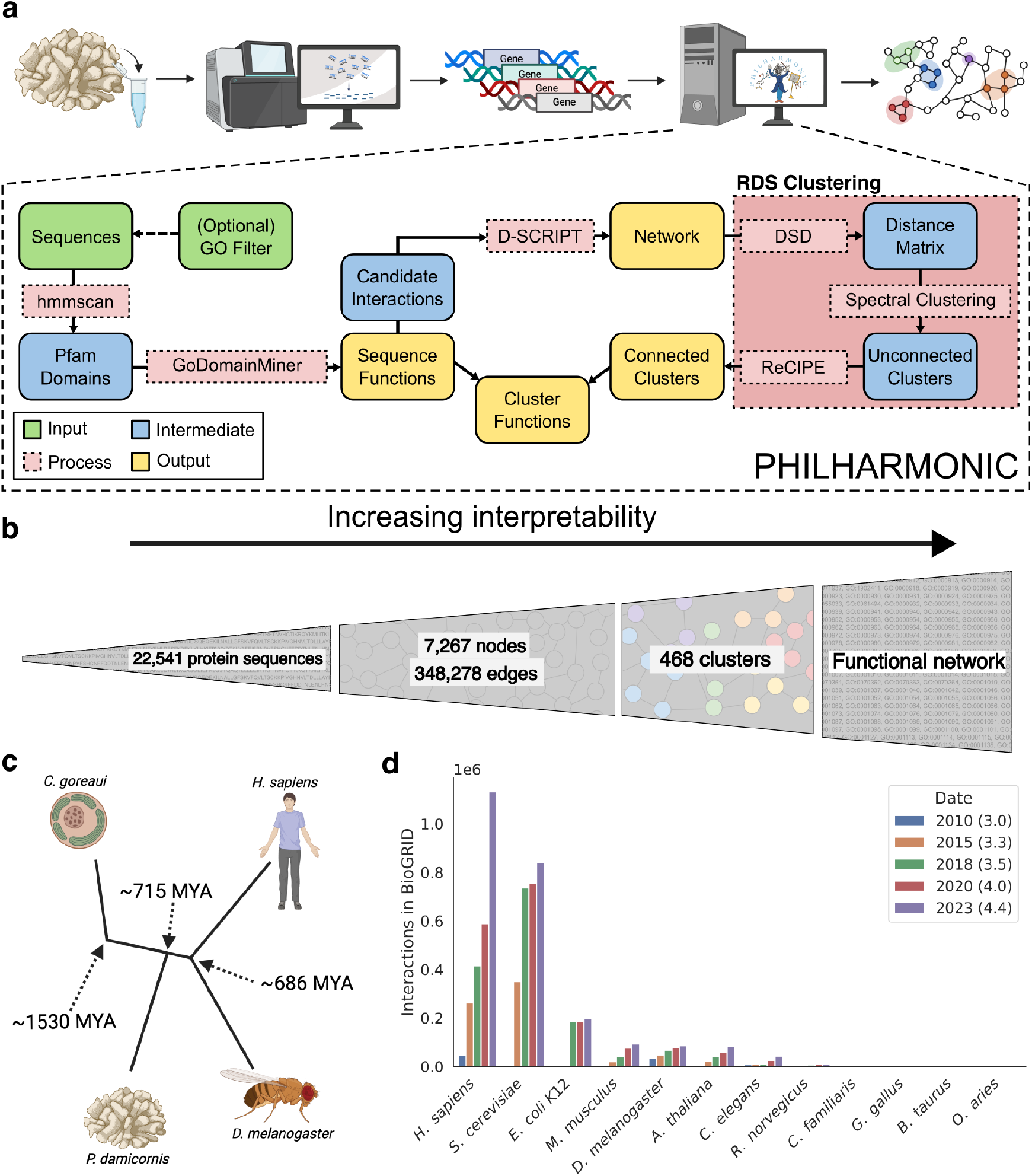
PHILHARMONIC is an engine for biological discovery in model and non-model organisms. **(a)** We address the problem of functional network inference in model and non-model organisms with PHILHARMONIC. Given a newly-sequenced proteome from a non-model organism, we use D-SCRIPT [17] to predict a *de novo* protein-protein interaction network (optionally on a smaller set of GO-filtered proteins, if computational cost is an issue). We cluster this network using RDS [22], which combines DSD [23] and hierarchical spectral clustering [19] with our newly developed ReCIPE method. We use hmmscan [24] and GoDomainMiner [25] to map sequence functions, then aggregate the *individual* sequence functions to assign *cluster* functions, which allows for network-level discovery and hypothesis generation. **(b)** PHILHARMONIC requires only protein sequences to run, which on their own provide little functional information. Starting with 11,487 *D. Melanogaster* or 22,541 *P. damicornis* sequences, we integrate several other sources of information, denoising, and downstream processing to build up to a much more interpretable functional network with meaningful communities. **(c)** For species like the cauliflower coral *P. damicornis* and its algal symbiont *C. goreaui*, hundreds of millions of years of evolutionary distance from well-studied model organisms make naive homology-based transfer of interactions or protein functions difficult. Even for well-studied model organisms like the fly *D. melanogaster*, a large evolutionary distance makes transfer from human data challenging. **(d)** We show the number of protein interactions in the BioGRID [6] database for several different organisms over several years (database versions). Nearly all known PPIs are concentrated in six well-studied model organisms, while most species have no known interaction networks.

There was likewise no existing computational method that could construct a sufficiently useful network. It is well-known that even over relatively short evolutionary distances there is substantial network re-wiring [8]. Even when sequence and protein structure conservation can allow confident ortholog assignment from a well-studied species, interaction patterns are often not preserved. The probability of shared edges decreases as the evolutionary distance between two species increases—so the networks from well-studied organisms such as human cannot trivially be used to transfer interactions to non-model organisms. Existing deep learning methods that predict physical PPIs directly from protein amino acid sequence are either too slow to make genome-wide predictions [9–12] or are not accurate enough to use the raw predicted network as ground truth interactions [13–15].

Here, we introduce **PHILHARMONIC**, the first method that successfully enables systems-level functional genomics in non-model organisms and accurately infers protein functions and communities genome-wide. With PHILHARMONIC, we annotate proteins that are too evolutionary distant from model organisms for existing methods to handle. Our key insight is that although protein-protein interaction networks predicted by high-throughput computational methods may not reach the level of experimental correctness, they nonetheless contain enough signal so that with innovative downstream processing, these *de novo* networks can serve as an information-rich scaffold, enabling pathway-level network analysis and functional genomics. We validate our method using experimental data and gold-standard functions in the fruit fly, and then show that PHILHARMONIC makes meaningful predictions of protein function and credibly sketches interacting modules in coral and algae. PHILHARMONIC provides annotations across hundreds of millions of years of evolution, shedding light on the “dark proteome,” [16] and bringing powerful systems biology approaches to non-model organisms.

## Results

### Overview of PHILHARMONIC

To overcome the lack of experimental functional data, we deploy lightweight, protein language model-based PPI prediction methods at the genome scale and build upon robust methods for remote homology functional inference. Then, our novel clustering and reconnection approach allows us to de-noise the predicted network, producing highly informative functional modules. This modular decomposition of the network then enables functional assignment at a community level, illuminating the function of proteins outside the reach of traditional homology-based methods.

PHILHARMONIC (Protein Human-transferred Interactome Learns Homology And Recapitulates Model Organism Network Interaction Clusters) requires only a set of protein sequences to perform *de novo* inference of functional communities (Figure 1a,b). First, we use the deep learning method D-SCRIPT [17] to infer the initial, noisy PPI network. We develop a novel hierarchical clustering algorithm based on the Double Spectral method, which couples a Diffusion State Distance (DSD)-based similarity metric [18] with several rounds of recursive spectral clustering [19] to convert the PPI network into putative functionally-enriched non-overlapping clusters of balanced sizes that are amenable to easy biological analysis (Methods). These clusters are by definition non-overlapping and can be highly disconnected, as opposed to in real biological systems where proteins often play many functional roles. To rectify this, we develop a novel method, ReCIPE (Reconnecting Clusters In Protein Embedding) that creates coherent, better connected, and more explainable clusters by re-adding high degree nodes with a greedy algorithm, thus optimizing intracluster connectivity. We show that adding ReCIPE on top of the DSD + Spectral non-overlapping clustering pipeline produces overlapping clusters that are superior by functional enrichment measures to existing state of the art unsupervised overlapping clustering methods on PPI networks. In this manuscript, we jointly refer to the clustering phase of Philharmonic (Recipe+DSD+Spectral) as RDS. We leverage robust remote homology methods that assign sequence-based gene ontology (GO) annotations to proteins based on Pfam domains [20], and our clusters inherit the functions that are enriched in its constituent proteins. On a gene-by-gene basis, we have previously demonstrated that some functional inference can still be accomplished by remote homology approaches using profile-profile HMMs [21]. We demonstrate here that these annotations can be successfully transferred to non-annotated proteins to infer larger functional pathways when combined with functionally-enriched clusters. Finally, we summarize the function of each cluster in paragraph form using generative AI to create easily readable summaries of cluster functions and to add biological interpretability. PHILHARMONIC provides a user-friendly Python package and Colab notebook that seamlessly runs the entire pipeline end-to-end, requiring only a sequenced proteome. We have designed PHILHARMONIC to be accessible to users even with limited programming expertise. Additional technical details are provided in Methods, and the full workflow is shown in Figure A1. PHILHARMONIC is available as a free and open-source tool that can be easily applied to a wide-variety of non-model organisms at https://github.com/samsledje/philharmonic.

### Validated cluster functions in *D. melanogaster*

Before exploring function in non-model organisms, we first sought to validate PHILHARMONIC within the fruit fly *D. melanogaster*. Due to efforts like the FlyBase project [26], the fruit fly is thoroughly well-characterized, and we have access to high-confidence functional annotations, experimentally validated protein interactions, and extensive gene expression data. These data allow us to demonstrate that PHILHARMONIC yields functionally meaningful communities in a “gold-standard” setting, contrary to non-model organisms such as coral where such data are sparse or non-existent.

We first use known GO function annotations from FlyBase [26], and define functional similarity between two proteins as the Jaccard similarity between the sets of GO [27] terms assigned to each protein. Then, we define cluster coherence as the mean functional similarity of all pairs of proteins in the cluster. We find significant functional coherence in PHILHARMONIC clusters (*p* = 5.2 × 10^−16^, one-tailed independent t-test) in the fruit fly (Figure 2a,b), compared to a random label assignment that preserves cluster size and connectivity. We construct this random assignment by shuffling node function assignments for the PHILHARMONIC clusters, preserving the degree and size distribution of clusters. We note that protein-level functions are *not* used for construction of either PHILHARMONIC or the random cluster baseline. We then replicate this analysis using FlyBase pathway membership to evaluate a higher level of functional relatedness (Appendix A.6.2, Figure A3), similarly finding that PHILHARMONIC clusters group proteins from shared pathways. In addition to coherent functions and pathways, we test whether genes in PHILHARMONIC clusters have correlated expression using dense time-series gene expression data in *D. melanogaster* from Schlamp et al. [28] (Appendix A.6.3). We show that PHILHARMONIC clusters are significantly more likely to have correlated expression than a random baseline (*p* = 4.79 × 10^−21^, one-tailed related t-test, Figure 2c,d). We then focus on the impact of D-SCRIPT for the network generation step, replicating the above GO functional analysis using the gold-standard *D. melanogaster* network from the STRING database [7] (Appendix A.6.1): this allows us to quantify the extent to which noise in the D-SCRIPT inferred network impacts functional inference. We find that inference using the noisy predicted network is consistent with that using the STRING network. We note that by validating in this setting, we remove the risk that cluster coherence arises solely from domain interaction bias in the predicted network.

**Figure 2:**
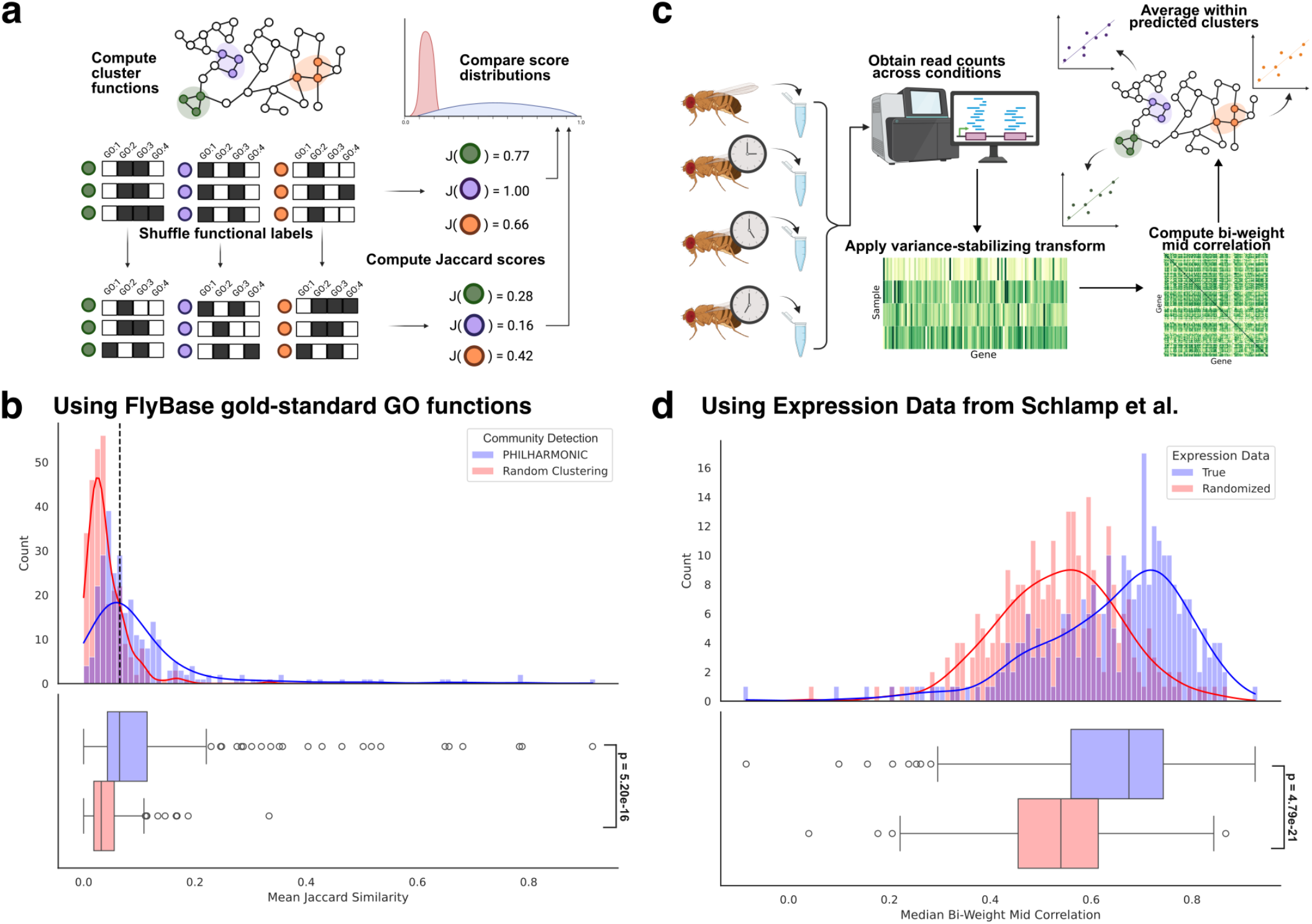
PHILHARMONIC clusters group functionally-related and co-expressed proteins in *D. melanogaster*. **(a)** We compute the functional coherence of a cluster as the mean Jaccard similarity between the sets of GO functional terms using gold-standard GO functions from FlyBase [26]. We compare these coherence scores to a distribution-preserving random clustering of the network by shuffling functional labels. **(b)** We find that PHILHARMONIC clusters contain proteins that are significantly more functionally related than the baseline (*p* = 5.20 × 10^−16^, one-tailed independent t-test). We set a threshold of 0.05 where the two curves cross, above which clusters are unlikely to be drawn from a random distribution. This can be used to select high-likelihood clusters, even in absence of gold standard functions like those from FlyBase. **(c)** We also evaluate clusters by evaluating whether genes within a cluster are co-expressed. Using time-course gene expression from Schlamp et al. [28], we first compute a variance-stabilizing transform on counts, then we compute the median bi-weight mid correlation between pairs of genes in the same cluster. We hypothesize that functionally related genes are more likely to be co-expressed across several conditions and samples. **(d)** We find that genes in PHILHARMONIC clusters are significantly likely to have correlated patterns of expression (*p* = 4.79 × 10^−21^, one-tailed related t-test) compared to a background re-shuffling of expression vectors. We emphasize that expression data was not used for clustering, only the patterns of network connectivity.

While our primary goal in applying PHILHARMONIC to the fly was to validate its performance with known functions and pathways, we show that even in a well-studied organism our approach can add new insight that could not be obtained by looking only at individual proteins or clusters. We analyzed the global organization of PHILHARMONIC clusters in the predicted *D. melanogaster* network. This network displays scale-free characteristics [29] typical of biological networks (Figure 3a), and PHILHARMONIC produces intentionally balanced cluster sizes (Figure 3b. We construct a graph representation of the network where each node is itself a cluster, which we use to investigate high-level functional organization (Figure 3c). We focus our further analysis on a cluster of 9 proteins (Figure 3d). This cluster was specifically selected based on the high-level organization seen in the cluster graph, as it is highly central in the graph and connects a diverse set of functions (a “bridge” cluster, betweenness centrality of 0.0291, 16th out of 285 clusters). We show all nodes that have at least 10 crossing edges to the bridge cluster. Proteins in this bridge cluster interact with proteins annotated for inflammatory response (orange, GO:0006954), mitotic cell cycle (green, GO:0000278), extracellular matrix, DNA replication (purple, GO:0006260), and DNA-templated transcription (light blue, GO:0006352) among others. In addition, this cluster is connected to several clusters which themselves contain proteins with diverse functional annotations (no one dominant function).

**Figure 3:**
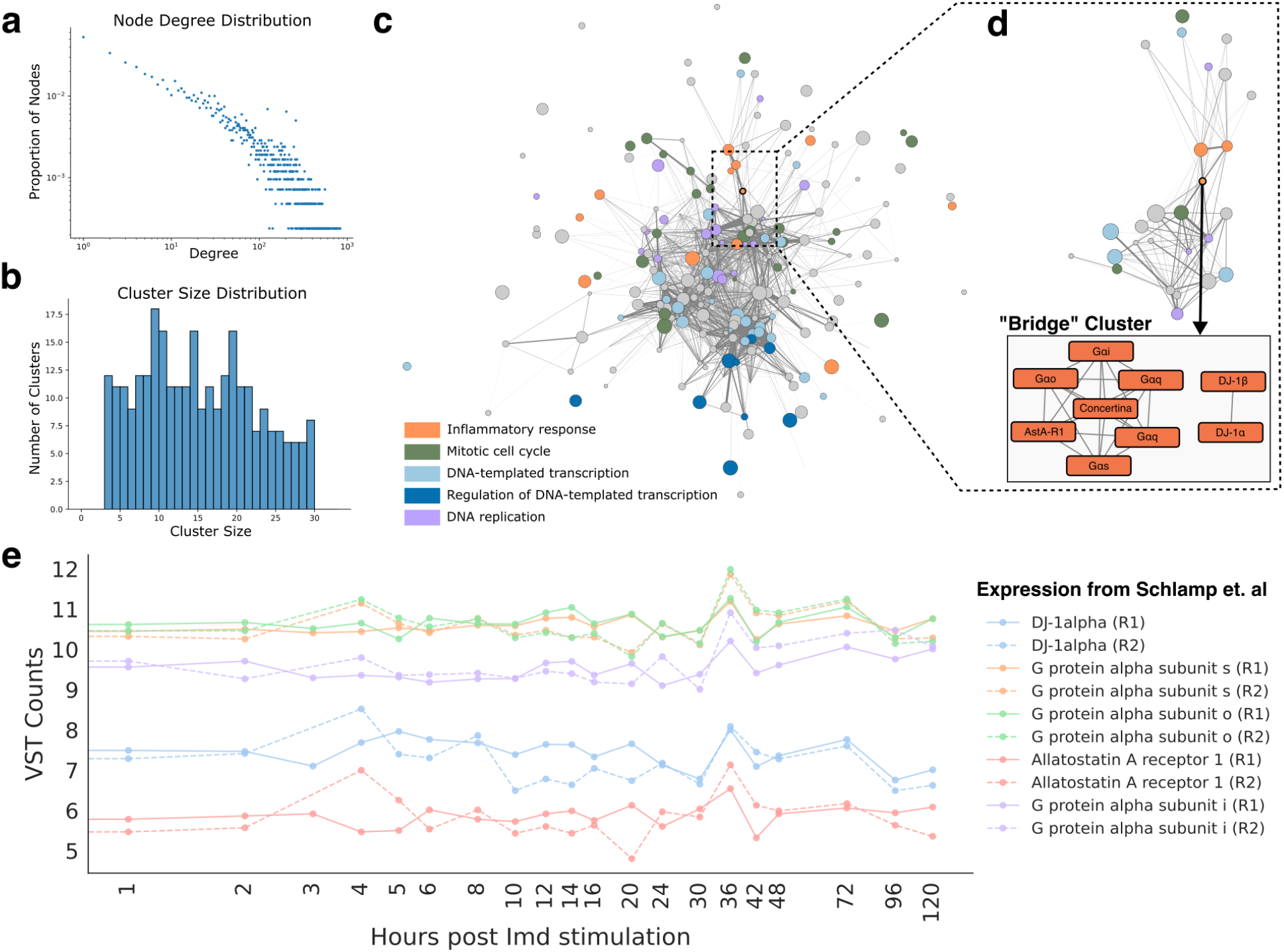
PHILHARMONIC recovers functional clusters in *D. melanogaster*. **(a)** The synthetic fly PPI network predicted by PHILHARMONIC follows a scale-free degree distribution commonly found in biological networks [29]. **(b)** PHILHARMONIC yields balanced clusters with sizes between three and 29 nodes. **(c)** We show a *“cluster graph”* for *D. melanogaster*, where each node is a cluster and edges indicate high cross-connectivity between clusters. For ease of visualization, we have filtered out clusters that do not connect to others with at least 50 crossing edges. **(d)** We identify a central “bridge” cluster that has one of the highest betweenness centralities in the cluster graph (Rank 16 out of 285 clusters; Betweeness centrality score: 0.0291). This cluster contains several G protein alpha subunits, as well as AstA-R1, DJ-1*α*, and DJ-1*β*, and is connected to a diverse set of functional clusters. We hypothesize that the proteins in this cluster play a crucial role in routing of environmental signals in *D. melanogaster*. **(e)** Members of the “bridge” cluster have correlated time-series expression after stimulation of the immune pathway [28].

This bridge cluster is primarily composed of G*α* subunits and the G*α*-like *concertina*/G*α*11/12. In mice, G*α*i and G*α*o proteins are known to be involved in hearing [30] and give differential response in response to food deprivation [31]. In the fruit fly, the G*α*s protein is involved in olfactory signal transduction [32] and G*α*q is known to play a role in photoreceptor signaling [33, 34] and the gustatory response [35]. Interestingly, this cluster also contains DJ-1*α* and DJ-1*β*, both of which have been shown to convey a neuroprotective role against oxidative stress in multiple contexts [36, 37]. There is some evidence that G*α*i and G*α*o are target proteins of reactive oxygen species [38], making this cluster a good starting place to begin the search for the mechanism by which this protective role is played. We hypothesize that through diverse interaction partners, the proteins in this cluster are processing environmental signals and routing them to the appropriate systems to respond, acting as conductors of cellular function. In addition to the literature, there is experimental support for this role; several proteins in this cluster have correlated expression time series in response to immune stimulation (Figure 3e).

### Dissecting the functional network of the coral *P. damicornis*

After validating PHILHARMONIC in the fly, we next sought to apply it in an important non-model system. Corals are a prime example of the type of non-model organism for which functional systems genomics has traditionally been difficult. Corals are key to the ocean’s vast biodiversity and provide significant economic, cultural, and scientific benefits [39–41]. While they cover only 0.1% of the ocean floor, coral reefs are home to the largest density of animals on earth, rivaling rain forest habitats in species diversity [42]. As a result of anthropogenic activities, coral holobionts are declining rapidly (Appendix A.2). This trajectory of loss highlights the extreme urgency to assist corals in overcoming the impacts of pollution and global warming before the damage to these vital reef ecosystems is irreparable. New efforts are underway to harness the growing amount of genomic information that is becoming available for the coral animal, its hosted symbiotic algae, and other elements of the coral holobiont, to discover ameliorative strategies that could slow coral decline [43]. However, these efforts are hampered by the great evolutionary distance of corals to any species with functionally well-annotated genomes (Figure 1c).

We first evaluated whether PHILHARMONIC could generate functionally meaningful clusters in the coral *P. damicornis*, measured by functional coherence. Here, because gold standard GO terms are not known, we define functional similarity between two proteins as the Jaccard similarity between the sets of GO [27] terms assigned to each protein by our Pfam-based function annotation. Then, the cluster coherence is the mean functional similarity of all pairs of proteins in the cluster. We find that *P. damicornis* clusters computed by PHILHARMONIC conserve significantly more functional relationships (*p* = 1.15 × 10^−53^, one-tailed independent samples t-test) than the baseline (Figure 4c). We also compare with several other state-of-the-art clustering methods (Appendix A.7.2, Table A3, Figure A5). Using the GO Slim term appearing most frequently assigned to proteins within that cluster), we evaluate the coherence of clusters with each high-level function (Figure 4e), and we identify several highly-coherent clusters related to mitotic cell cycle (GO:0000278, green), DNA-templated transcription (GO:0006351, light blue) and its regulation (GO:0006355, dark blue), and inflammatory response (GO:0006954, orange). We perform the same analysis using only GO Slim [27] terms (Appendix A.7.1, Figure A4) and we replicate this analysis on the *C. goreaui* network (Appendix A.8).

**Figure 4:**
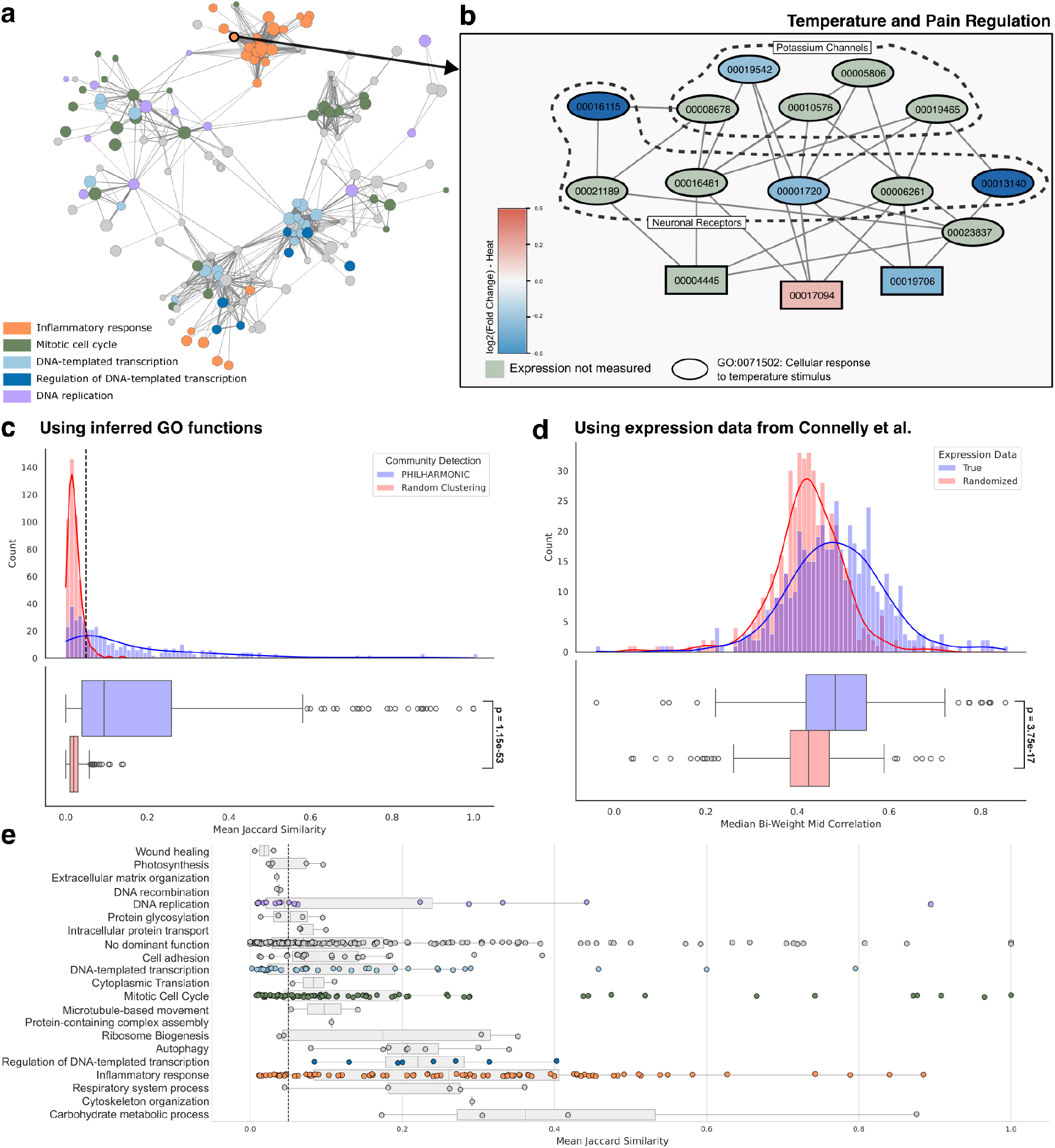
*De novo* network inference and functional clustering in *P. damicornis*. **(a)** The *P. damicornis* cluster graph, where nodes are clusters and edges are drawn if *>*= 50 PPIs exist between proteins in each cluster. We highlight here five high-level cluster functions. We find neighborhoods of the network focused on inflammatory response (GO:0006954, orange), mitotic cell cycle (GO:0000278, green), and DNA-templated transcription (GO:0006351, light blue). Clusters focused on regulation of transcription (GO:0006355, dark blue) and DNA replication (GO:0006260) are more dispersed throughout the network (Appendix A.7.5). **(b)** We highlight a cluster involved in temperature and pain regulation in *P. damicornis*. Here, each node is a protein and an edge is a predicted PPI. For legibility, we have dropped the “pdam “ prefix from protein identifiers on node labels. We find that 12 of the 15 proteins in this cluster are annotated with cellular response to temperature stimulus (GO:0071502) (indicated with ovals). Nodes are colored by *log*_2_(Fold change) in expression when exposed to heat [44] (genes not in data are colored green). This cluster contains several putative potassium channels and neuronal receptors. **(c)** PHILHARMONIC clusters are significantly more functionally coherent (*p* = 1.15 × 10^−53^ one-tailed independent t-test) than would be expected by chance, computed as the median Jaccard score between GO terms assigned to each pair of proteins in the same cluster. We set a cluster coherence threshold of 0.05 (dashed grey line) where the curves cross. **(d)** We find that proteins that share a PHILHARMONIC cluster are significantly more likely to be co-expressed than a baseline with re-shuffled expression data (*p* = 3.75 × 10^−17^, one-tailed related t-test). **(e)** We show each of the 468 clusters, separated by GO Slim functional assignment and plotted by cluster coherence. We find a substantial number of clusters above the 0.05 threshold across several diverse functions, including those highlighted in the cluster graph above.

We next investigated whether the PHILHARMONIC *P. damicornis* clusters are more likely to group proteins which are co-expressed. We use gene expression data from Connelly et al. [44], consisting of 47 samples of RNAseq data from four *Pocillopora* coral specimens, collected in triplicate across four conditions (control, heat exposure, antibiotics exposure, heat+antibiotic exposure). We find that proteins in PHILHARMONIC clusters are significantly more likely to have correlated expression across these tested conditions (*p* = 1.27 × 10^−21^, one-tailed related samples t-test) (Figure 4d). We also compare against a clustering derived directly from functional labels, and find similar levels of co-expression, but with significantly better connectivity in PHILHARMONIC clusters (Appendix A.7.3, Figure A.7.3). We compute an alternate method of co-expression within a cluster using the singular values of the cluster sub-matrix with similar findings (Appendix A.7.4, Figure A7).

We find substantial organization within the *P. damicornis* cluster network (Figure 4a), with different functions localized to different “neighborhoods” of the network. This view of the network highlights the centrality of transcription, gene regulation, and the cell cycle in cellular function [45], and we identify tightly-connected neighborhoods of the overall network dedicated to inflammatory response [46], mitotic cell cycle, and DNA-templated transcription (Appendix A.7.5, Table A5). We perform a similar high-level analysis of the *C. goreaui* network, including two additional case studies (Appendix A.8).

### Clustering identifies the regulation of thermal response

In addition to this global view, we investigated two clusters in depth, both of which involve ion channels communicating with *G*-coupled protein receptors (GPCRs). We analyze here one PHILHARMONIC cluster involved in cellular response to temperature stimulus (GO:0071502) (Figure 4b). Cnidarians, including coral animals, have a large and diverse family of voltage-gated potassium ion (K^+^) channels, but little is known about how the function of these channels differs across different cnidarian species [47]. This cluster includes three proteins that are likely voltage-gated K^+^ channels (`pdam_00019542`, `pdam_00019465`, `pdam_00010576`), a cyclic nucleotide-gated K^+^ channel (`pdam_0008678`, [48]), and a hyperpolarization-activated cyclic nucleotide-gated (HCN) channel (`pdam_00005806`). HCN channels are involved in controlling neuronal excitability, dendritic integration of synaptic potentials, synaptic transmission, and rhythmic oscillatory activity in individual neurons and neuronal networks in humans, where they have been studied for their role in epilepsy and pain [49]. Cnidarian HCN channels share all major functional features of vertebrate HCNs, including reversed voltage-dependence, activation by cAMP and PIP2, and blocking by extracellular cesium ions [50]. There is some evidence that HCN channels are also involved in rhythmic firing and the so-called pacemaker channels [51], and this cluster also contains several GPCRs that are predicted to bind to and likely modulate the functions of the above diverse ion channels in response to different stimuli. Two of these, `pdam_00016481` and `pdam_00016115`, are similar to *ADRB1* and *ADRB2* respectively, which are involved in brown adipose tissue differentiation and linked to heat generation in humans [52]. Notably, many of these receptors are linked to neural peptides (Appendix A.7.6), including an orexin receptor (`pdam_00006261`) that has been implicated in systems involving sleep and light-sensing in mammals [53]. `Pdam_00001720` resembles the allatostatin-A receptor, to which the neuropeptide allatostatin-A binds, modulating feeding behavior in mosquitos [54]. We identify one protein in this cluster that is poorly characterized by sequence homology (`pdam_00017094`), though structural homology searches using Foldseek show a conserved structural domain similar to Alpha-(1,6)-fucosyltransferase. Using data from Connelly et al. [44], we find that `pdam_00017094` is over-expressed during heat stress. We predict this protein to interact with a subset of the GPCRs, including the putative orexin and allatostatin-A receptors. We investigate a separate case study cluster related to temperature response in Appendix A.7.8.

### Uncovering the neurophysiological response to environmental stimuli

We compared our RDS community construction approach (Figure 5a) with several other popular clustering methods, and we find that not only does our approach result in more balanced cluster sizes (Table A4) and interpretable clusters, but that reconnection with ReCIPE actually further increases functional coherence of clusters (Figure 5b, Appendix A.7.2). Following this insight, we next highlight a cluster that has been strongly re-connected by ReCIPE, one enriched for putative neurological receptors and involved in the coral response to environmental stimuli (Figure 5c, Appendix A.7.7). This cluster contains two proteins with high levels of homology to K^+^ voltage-gated channels [55], namely `pdam_00001381` in the original cluster and `pdam_00006320` among the proteins added in by ReCIPE. Among the remaining proteins include several that are highly homologous to ligand-gated ion channels, previously recognized as key modifiers of behavior in vertebrate brains. We also identify four gamma-aminobutyric acid (GABA) receptors (`pdam_00012376`, `pdam_00001388`, `pdam_00001389` and `pdam_00024051`, the last added by ReCIPE), and four proteins that are homologous to nicotinic receptors in vertebrates (`pdam_00006972`, `pdam_00015999`, `pdam_00006973`, `pdam_00012507`). GABA has been identified as a signaling molecule in feeding behavior, orientation, and tentacle movement in other Cnidarian species [56]. Jing et al. showed that the roles of GABAergic inhibitory interneurons can also be extended to the *Aplysia* feeding motor network [57]. They are widely expressed in the central nervous system, while, in the periphery, they mediate synaptic transmission at the neuromuscular junction and ganglia. There are three proteins in this cluster which are poorly characterized by sequence homology—`pdam_00011550`, `pdam_00018139`, and `pdam_00006109`. Two of these, `00011550` and `00006109`, are likely interacting with a GABA receptor candidate `00024051` that is strongly under-expressed in heat conditions. We hypothesize that proteins in this cluster are involved in the coral’s response to its environment, acting as a sort of nervous system. Because this hypothesis rests on the organization of the cluster rather than any individual protein function, only our network-based approach could have arrived at this hypothesis.

**Figure 5:**
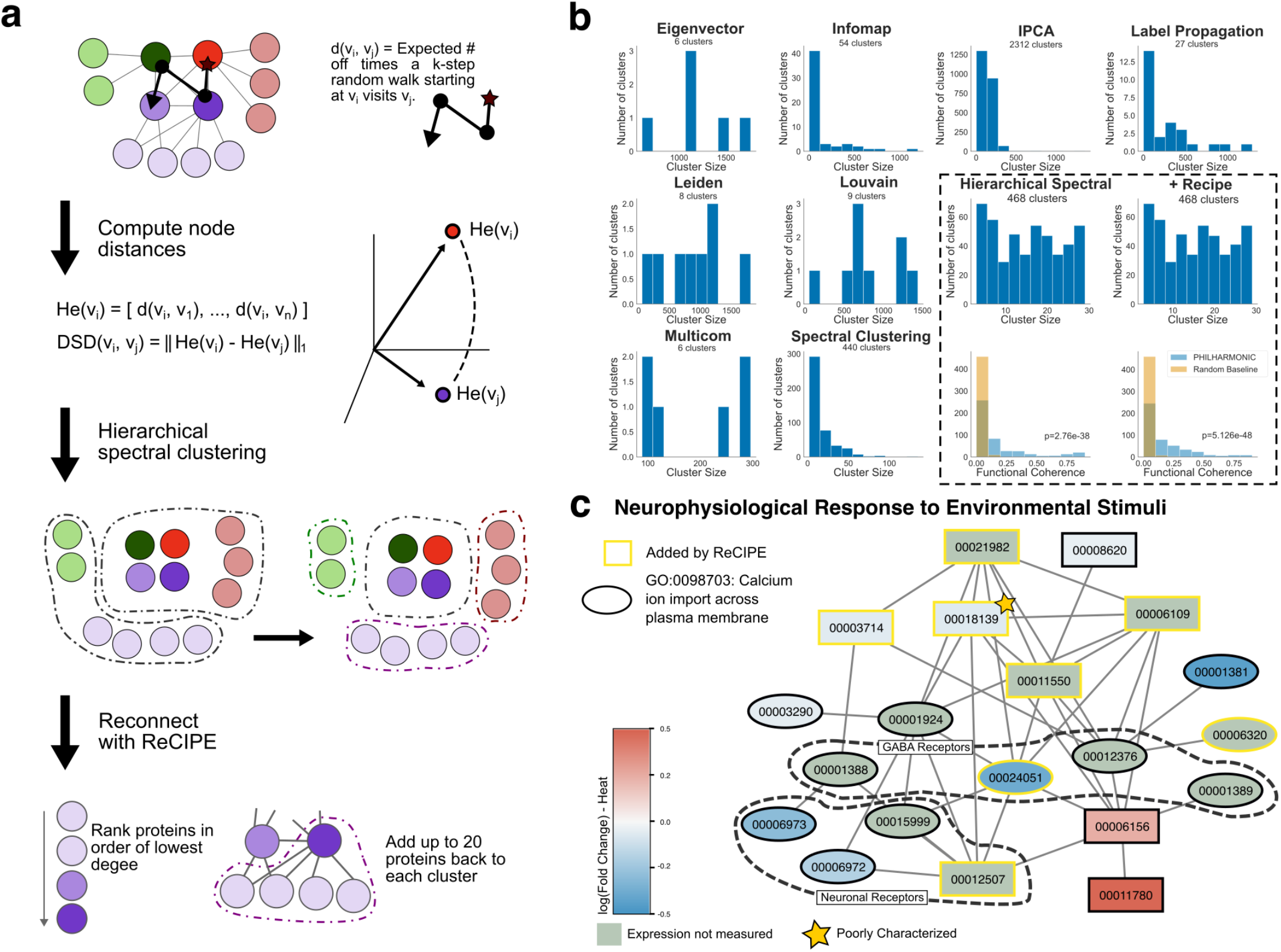
Reconnecting with ReCIPE increases interpretability and functional coherence of clusters. **(a)** Schematic overview of the RDS clustering pipeline. DSD [18] defines pairwise node distances based on the expected distribution of random walks on the graph. We recursively perform spectral clustering on this distance matrix until a cluster size threshold is met. However, this often results in clusters that are difficult to interpret, because there are missing hubs that would connect the proteins. To address this, we use ReCIPE to identify candidate nodes for re-addition, rank them prioritizing low-degree nodes, and add them back greedily until a connectivity threshold or maximum number of proteins is hit. **(b)** We compare our clustering pipeline with several other popular clustering approaches (using CDlib [58]). Our approach achieves much more balanced and interpretable cluster sizes than these alternatives, as well as significantly better functional coherence (Appendix A.7.2). **(c)** We show another case study, a cluster involved in neurophysiological response to environmental stimuli in *P. damicornis*, where ReCIPE has re-added several important proteins to the cluster. Proteins annotated with calcium ion import across the plasma membrane (GO:0098703) are indicated with ovals. Nodes are colored by *log*_2_(Fold change) in expression when exposed to heat [44] (genes not in data are colored green). This cluster contains several putative GABA and neuronal receptors.

## Discussion

Here, we developed PHILHARMONIC to enable inference of protein interactions and functions across the tree of life. Our approach combines protein language models, spectral algorithms, and traditional HMM-based sequence algorithms to generate functionally meaningful and biologically interpretable clusters. We validate the quality of our inferred networks and clusters with substantial support from functional coherence, gene expression, and the literature. These results demonstrate the power of applying a network lens to function, by identifying hierarchies of organization and propagating function throughout the genome. In *D. melanogaster*, gold-standard functional data and independent gene expression data sets validate that PHILHARMONIC generates high-quality and functionally meaningful clusters.

Further, we show that these results extend beyond a single species and apply PHILHARMONIC to characterize the coral *P. damicornis* and its algal symbiont *C. goreaui*. Using PHILHARMONIC, we identify clusters related to thermal and environmental response in the coral *P. damicornis*, providing new insight into coral resilience to rising ocean temperatures. We also identify clusters in the symbiont related to detoxification and the response to oxidative stress. Non-model organisms such as these play a crucial role in our world, from the impact of coral on reef ecosystems [41], to the value of livestock and crops in agriculture, to the potential of microbes to combat climate change through carbon capture [59] or digestion of plastics [60]. Recent research suggests that certain biological mechanisms in these organisms may actually be better conserved with respect to human biology than those from the traditional model organisms [61]. While substantial previous work has advanced our understanding of functional networks in humans [62–66], limited experimental data and the large evolutionary distance from well-studied organisms has made similarly in-depth analysis of functional networks in non-model organisms difficult. PHILHARMONIC presents a meaningful step in improving functional coverage for these important organisms. Even in better-studied organisms, PHILHARMONIC provides value in building confidence in functional annotations. While a function derived from a single source may be lower confidence, multiple occurrences of the same function in a cluster is a strong signal of the validity of that function.

As researchers work to decode the genomes of diverse organisms [67–69], PHILHARMONIC enables a network approach to understanding protein function, but the results must be carefully interpreted. We note that assessment of protein function prediction is itself a challenging and well-studied problem [70], and that our current evaluation scheme bears some risk of circularity. If predicted edges are biased towards proteins with similar domains, then the clusters from this network are more likely to be functionally enriched simply due to this domain similarity. While we have attempted to rule this out by evaluating coherence on gold standard networks as well, some risk still remains. In addition, PHILHARMONIC will sometimes produce clusters that give no functional insight due to limited functional transfer over extremely long evolutionary distances, errors in the underlying network, or spurious clustering of proteins (Appendix A.11). We also note that the components of PHILHARMONIC are modular, and as individual components improve, the overall workflow can as well—for example, D-SCRIPT could be replaced by improvements in PPI prediction [71], the Pfam search could be replaced by improvements in finding remote orthologous genes [72], or recent work on direct function prediction using protein langauge could be applied [73]. Despite these current limitations, the broad applicability and ease of use of PHILHARMONIC will enable domain experts to organize the proteome into functional clusters, to identify testable functional hypotheses, and to dissect and understand the complex cellular systems of these organisms.

## Acknowledgments

This work was supported by the US National Science Foundation under the “Harnessing the Data Revolution” NSF Big Idea: Grants No. 1939249, No. 1940169, No. 1939699, No. 1939795, and No. 1939263. S.S. was supported by the NSF Graduate Research Fellowship under Grant No. 2141064 and by the Simons Foundation. K.D. and L.C. were additionally supported by NSF No. 1934553. C.V. thanks NSF DIAMONDS REU No. 2149871. B.B. is supported by the NIH NIGMS 1R35GM141861. L.C. is supported by the National Institute Of General Medical Sciences of the National Institutes of Health under Award Number R01GM163241. The content is the sole responsibility of the authors and does not necessarily reflect the views of the NIH. We thank Chris Edelmaier, Zhicheng Pan, Natalie Sauerwald, and Bargeen Turzo for helpful discussions and feedback on the manuscript, as well as the anonymous referees for helpful feedback on the manuscript.

## Materials and Methods

### Data Collection

In this work, we focus on three species: the fly *Drosophila melanogaster*, the coral *Pocillopora damicornis*, and its dinoflagellate symbiont *Cladocopium goreaui* (formerly named *Symbiodinium goreaui*, Clade C type C1). Fly protein sequences were downloaded from FlyBase [26]. After filtering to proteins with between 50 and 800 residues due GPU VRAM limitations, we keep 11,487 fly sequences. Gold standard pathways and interaction networks for benchmarking were also downloaded from FlyBase. Coral protein sequences were obtained from Cunning et al. [74], while symbiont sequences were obtained from ReefGenomics [75, 76], based on Liu et al. [77]. After length filtering, we were left with 22,541 *P. damicornis* sequences and 28,271 *C. goreaui* sequences.

### Functional Annotation of Individual Genes

Remote homology methods can find related genes that might preserve functional roles over greater evolutionary distance than standard sequence-based or ordinary-HMM-based searches. We employ a two-step functional annotation to maximize conserved function. We construct an HMM database from Pfam [20] domains using hmmer [24], then use hmmscan to search our input sequences against this database. Each protein is then annotated with some number of domains. Finally, we use mappings between domains and GO biological process (GO:BP) functions from GODomainMiner [25] to transfer functional terms to candidate sequences. Because structured domains are more likely to be conserved across long evolutionary distance, this procedure ensures the highest quality functional annotations for single proteins—but still results in some proteins which are unable to be functionally characterized, or may yield false positive annotations, necessitating a systems approach to function.

### Synthetic Network Prediction

D-SCRIPT [17] is a deep learning method that was trained on human PPIs and has been shown to generalize broadly across species. It requires only the amino acid sequences of two proteins and predicts a probability of interaction. D-SCRIPT can either be run on all pairs of proteins, or, if there are computational constraints, instead the set of proteins can be curated based on a list of GO terms. In our case, we use a hand-curated list of diverse high-level GO:BP terms potentially related to coral metabolism, resilience, and regulation (Appendix A.4, Table A1). We keep any proteins that are annotated with these high-level GO terms or any child GO terms. We generate a list of candidate interactions by considering all pairwise interactions between the filtered set of proteins, resulting in 45.9 million coral candidates, 52.7 million symbiont candidates, and 15.2 million fly candidates. We use D-SCRIPT because it is both highly accurate for non-model organisms and computationally efficient enough to be run genome-wide on the proteins in a new organism—it takes approximately a week to predict ≈ 50 million interactions on a single GPU. We consider a positive interaction to be anything predicted with a probability greater than 0.5.

### Diffusion State Distance

In this manuscript, we develop a clustering pipeline consisting of a distance metric computation (DSD), a community detection step on those distances (Hierarchical Spectral Clustering), and a cluster reconnection step (ReCIPE). In this manuscript, we refer to the full clustering pipeline of DSD + Hierarchical Spectral Clustering + ReCIPE as “RDS”. We first describe DSD. DSD is an effective way to measure distance between nodes in a PPI network. We first consider the case where the edges are unweighted as in [18]. For two nodes (proteins) *a* and *b*, we measure the expected number of times a *k*-step random walk starting at *a* visits *b* (including the 0-step random walk), and call this *He*^*k*^(*a, b*). If *v*_1_, … *v*_*n*_ represent all the nodes in the network, we then associate to each node *v*_*i*_ the vector consisting of these *He*^*k*^ values to all *n* nodes, namely

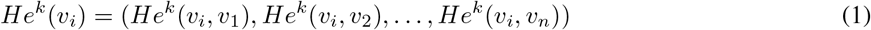

Then the DSD distance between two nodes *v*_*i*_ and *v*_*j*_ is defined as the *L*_1_ norm of the vectors *He*^*k*^(*v*_*i*_) and *He*^*k*^(*v*_*j*_), i.e.

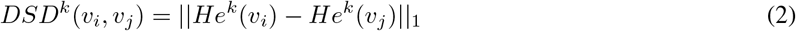

Cao et al. (2013) [18] show that this converges (rapidly, in small world networks) as the length of the random walk *k* → ∞, and denote the *DSD*(*v*_*i*_, *v*_*j*_) without the *k* superscript to mean the converged DSD. This can be computed exactly with the closed form in [18]. Furthermore, when there are weights on the edges, a straightforward modification presented in Cao et al. (2014) [23] can take into account confidence weights on the edges by instead biasing the underlying random walk to more likely go along edges of more confidence (in direct proportion to the confidence ratios among the edges from the node). Everything else in the weighted DSD definition stays the same. We set the confidences according to the edge weights predicted by D-SCRIPT.

### Community Detection by Hierarchical Spectral Clustering

Given the DSD-smoothed node distance matrices, we convert this to a similarity measure between nodes using the RBF kernel [78]. We cluster this distance matrix using a modified version of the spectral clustering algorithm described in [19], recursively splitting any cluster that is bigger than assigned threshold. The original DSD + spectral clustering method won the disease module identification challenge in DREAM 2016 [1]. Our updated version of this algorithm, which we call “Hierarchical Spectral,” gives us more control over the final distribution of cluster sizes using three parameters *K, D, M*. For the default PHILHARMONIC implementation, we initially call spectral clustering with *K* = 500 clusters. The initial set of clusters is recursively split, where clusters are recursively divided into 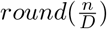 clusters (where *n* is the size of the cluster) until 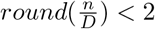. Since the network will display cluster organization at multiple scales, we chose the default *D* = 20 to create clusters of a size reasonable for biologists to further investigate by hand (resulting in a maximum cluster size of (1.5 × *D*) − 1 = 29). We finally discard clusters with fewer than *M* = 3 nodes as uninformative.

### Reconnecting Clusters with ReCIPE

We observed that because DSD groups together nodes that are most similar, the Hierarchical Spectral Clustering step occasionally placed nodes in the same cluster because they were all connected to the same intermediate-degree minor hub nodes, but these hub nodes were instead grouped with other minor hub nodes, producing highly disconnected clusters. We develop a new method, ReCIPE (Reconnecting Clusters in Protein Embeddings) to return the minor hubs back into these clusters if they were otherwise disconnected, allowing some overlap between clusters.

ReCIPE is a greedy algorithm that aims to reconnect disconnected clusters by selectively reintroducing proteins that were excluded during the initial clustering process. Proteins were considered for re-inclusion in the cluster (“candidate proteins”) if they connected at least *r* = 0.1 of the disconnected cluster components. Candidate proteins are added back in increasing order of degree. If *n* is the number of proteins in the cluster, and *c* is the number of disjoint cluster components, then proteins are added until the connectivity score 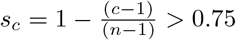, up to a maximum of *m* = 20 re-added proteins (where we note that a fully connected cluster has a *s*_*c*_ of 1, and that since clusters contain at least 3 nodes, the denominator is always defined (and at least 2)). We provide a full description of ReCIPE in Algorithm 1, and we provide pseudocode in Appendix A.9. ReCIPE is a substantial methodological innovation, because by relaxing the strictly non-overlapping requirements of spectral methods we maintain the strengths of those methods while gaining the flexibility and interpretability required to model complex biological systems.

We perform a comprehensive evaluation of hyperparameters both for community detection and reconnection in coral (Appendix A.10.1), yeast (Appendix A.10.2, Figure A11), and human (Appendix A.10.3, Figure A12) networks, and find that RDS is robust to a wide variety of choices for these parameters. We also show that the RDS method does not result in clusters that are massively overlapping and that even after the ReCIPE reconnection step, most nodes are not placed in too many clusters (Figure A14).

#### Algorithm 1 ReCIPE cluster reconnection algorithm

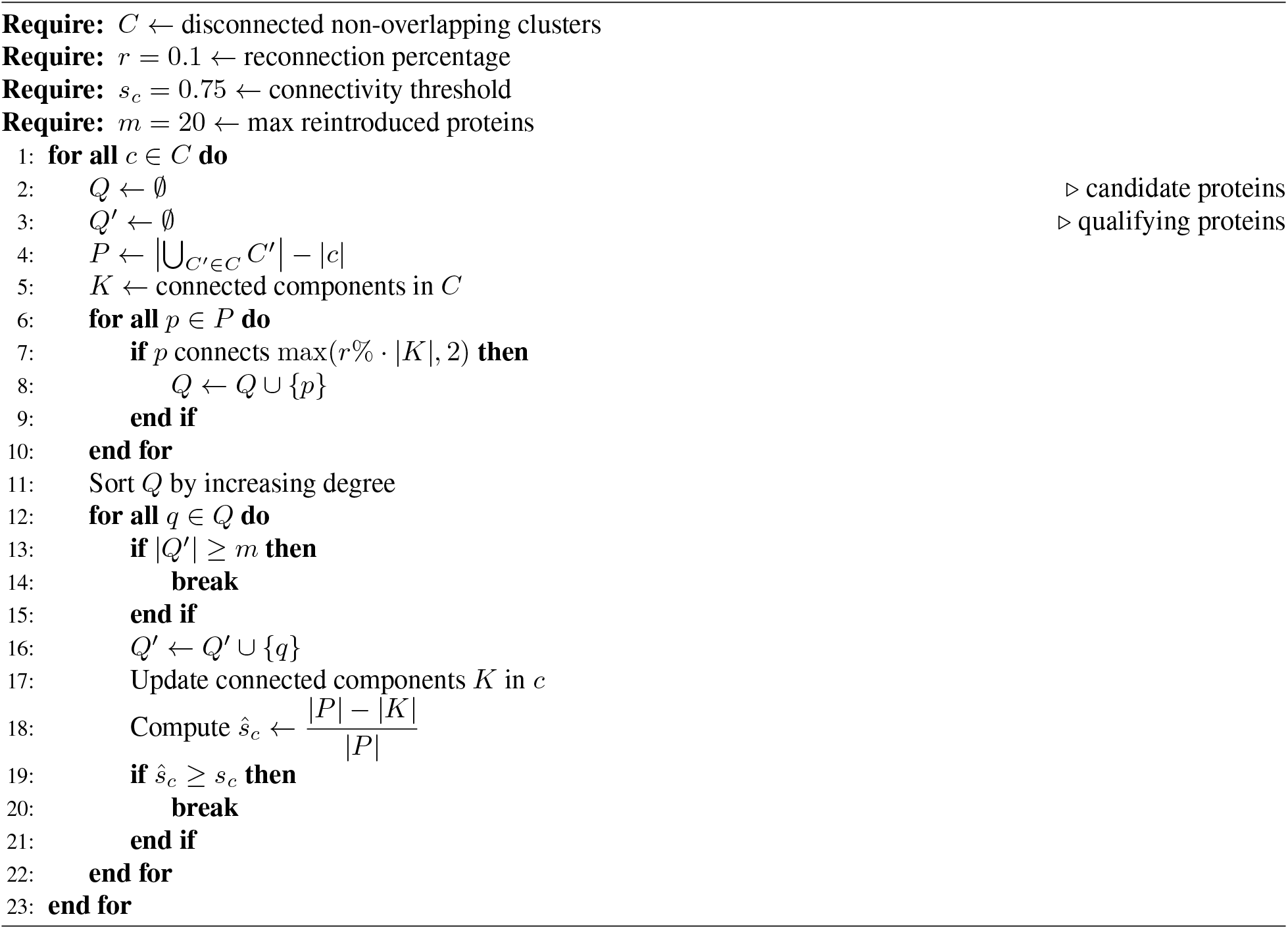

### Creating the Cluster Graph

To aid in the analysis of functional clustering in the network, we compute a higher-order graph where each node is not a protein, but rather a PHILHARMONIC-determined cluster. Given cluster *C* with nodes {*c*_1_, …, *c*_*n*_}, cluster *D* with nodes {*d*_1_, …, *d*_*m*_}, and original adjacency matrix *A* ∈ {0, 1}^*N* ×*N*^,*N > n, m*, we construct a new graph *G* with edges 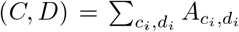. Finally, we drop all edges with weight less than *t* = 10, although we later filter to higher levels of *t* for visualization. When constructing this graph, we do not consider ReCIPE-added nodes in the edge count, as ReCIPE results in potentially overlapping clusters, leading to potentially double-counting edges if the same node appears in both *C* and *D*. We assign each cluster node a single function, based on GO Slim [27] terms appearing most frequently within that cluster. A given GO term must appear in at least 3 proteins in the cluster to be assigned, otherwise the cluster is left with an unassigned function.

### LLM Summarization

Inspired by work from Hu et al. [79], we sought to improve the interpretability of our clusters by using large language models (LLMs) to automatically annotate them with names. Not only does this more easily summarize cluster function, but we found that it eases communication within teams of users when referring to clusters. We utilize the `llm` command line utility [80], prompting the GPT 4o-mini model with a short text summary of each cluster containing the size, number of edges/triangles, and the top-represented GO terms within each cluster. The model is instructed to act as an expert biologist, and to come up with a short, descriptive, human-readable name for each cluster. The full system prompt is included in Appendix A.5. This name is added to the text summary from above and output for biological analysis. We note that, similarly to Hu et al., the correctness of cluster labels will be highly dependent on the prompt and the choice of LLM. These cluster names should be used as a helpful supplement rather than an absolute statement of truth about the function of the cluster.

### Implementation

We implement PHILHARMONIC using Snakemake [81]. This allows for consistent and reproducible analysis, flexible configuration with a `.yaml` file, and flexible allocation of computational resources. We use HMMER v3.3.0, D-SCRIPT v0.2.8, and fastDSD v1.0.1. ReCIPE is a newly implemented and `pip`-installable Python package; we use v0.0.4 in this work. We use the spectral clustering implementation from scikit-learn version 1.5.0. Statistical tests were performed with scipy version 1.11.3. We use Python version 3.11 and Snakemake version 8.10. We use version 0.16 of the llm command line utility. D-SCRIPT was run on a single A6000 NVIDIA GPU, and hmmscan was run using 32 cores. Cytoscape 3.10.2 was used for visualization of all networks, both protein-protein interaction and higher level cluster graphs. We provide a set of Cytoscape style files that can be used to visualize assigned cluster functions. PHILHARMONIC itself is available as a `pip`-installable package. Each clustering and analysis step is implemented as a Python script which is run by Snakemake, and can be individually called through the PHILHARMONIC package. We provide job submission scripts for running on HPC environments, and we also provide Google Colab notebooks to easily run PHILHARMONIC in the browser.

## Data Availability

Proteome sequences, predicted networks, clusters, and functional annotations for *P. damicornis, C. goreaui*, and *D. melanogaster* have been deposited on Zenodo at https://zenodo.org/records/15066518.

## Code Availability

PHILHARMONIC is available at https://github.com/samsledje/philharmonic. ReCIPE is available at https://github.com/focitti/ReCIPE.

