## Supplementary Material for "Decoding the Functional Interactome of Non-Model Organisms with PHILHARMONIC"

### A Appendix

#### A.1 Counts of PPI in different species

We constructed Figure 1c by downloading the specified versions of the BioGRID [6] database from `https://downloads.thebiogrid.org/BioGRID` and grouping interactions by species. We find substantial growth in the number of known human and yeast PPIs in the specified time frame, but relatively much less growth in even other model organisms, let alone non-model organisms.

#### A.2 The coral holobiont

Coral colonies are comprised of clonal cnidarian polyps that depend on a symbiotic relationship with algae in the family Symbiodiniaceae [82]. These dinoflagellate algae harvest light and synthesize nutrients in exchange for habitat and nitrogen sources [41]. Originally thought to primarily include endosymbiotic algae, the symbiosis is now known to extend to a much more complex community. The collective of thousands of bacteria, bacteriophages, viruses and fungi, in addition to Symbiodiniaceae and cnidarian, is known as the coral *holobiont* [83, 84]. Mass coral bleaching, or the expulsion of the symbiotic algae due primarily to thermal stress driven by marine heatwaves, is resulting in substantial coral mortality [85]. A recent study assessed 100 worldwide locations and found that the annual risk of coral bleaching has increased from an expected 8% of locations in the early 1980s, to 31% in 2016 [85], and this was before some of the most acute heat waves of the early 2020s.

#### A.3 Implementation of PHILHARMONIC

We implement PHILHARMONIC using the Snakemake package. Figure A1 shows the entire pipeline, generated using the command `snakemake --configfile config.yml --filegraph | dot -Tpng > img/pipeline.png`. This full figure shows not only the primary steps described in the main text, but additional necessary steps such as downloading required databases, preparing the hmm database, initial filtering and generating candidates for PPI prediction, and compiling all results together.

#### A.4 List of GO Slim terms for initial filtering

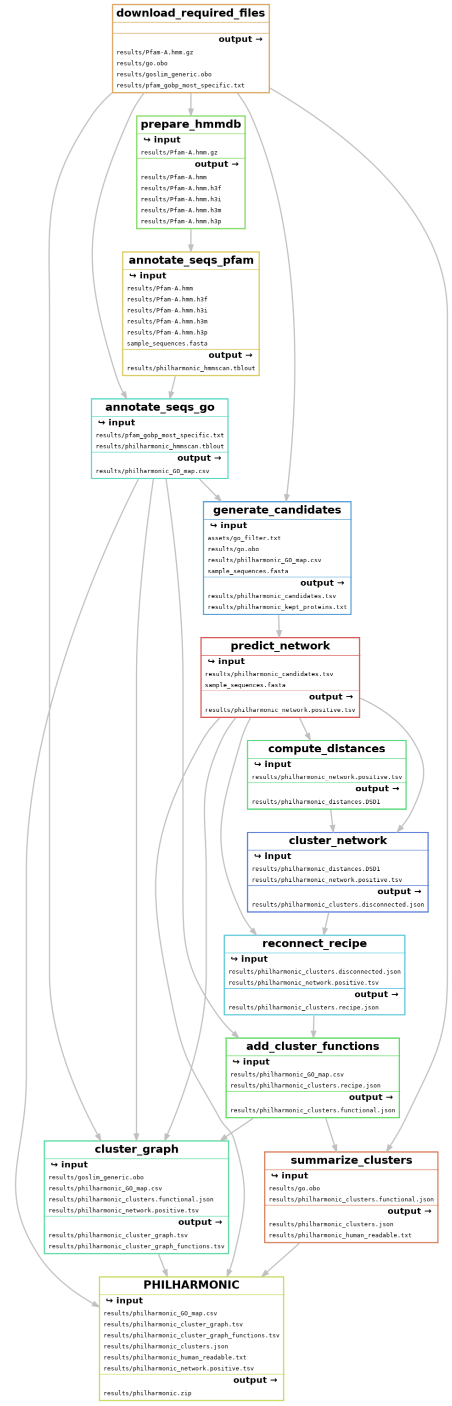


**Figure A1: Full PHILHARMONIC implemented in Snakemake**

**Table A1: GO Filter List.** We select a subset of high-level GO Slim terms, and filter candidate proteins to any annotated with these terms and any of their children. This allows us to focus on a functionally interesting subset of the network. We allow the user to provide their own list of GO filters, or to not filter at all.

| GO Term | Description |
|---:|---:|
| GO:0098754 | detoxification |
| GO:0098542 | defense response to other organism |
| GO:0050886 | endocrine process |
| GO:0050877 | nervous system process |
| GO:0048870 | cell motility |
| GO:0042060 | wound healing |
| GO:0023052 | signaling |
| GO:0022600 | digestive system process |
| GO:0015979 | photosynthesis |
| GO:0012501 | programmed cell death |
| GO:0007018 | microtubule-based movement |
| GO:0006954 | inflammatory response |
| GO:0006914 | autophagy |
| GO:0006790 | sulfur compound metabolic process |
| GO:0006766 | vitamin metabolic process |
| GO:0006629 | lipid metabolic process |
| GO:0006575 | cellular modified amino acid metabolic process |
| GO:0006520 | amino acid metabolic process |
| GO:0006486 | protein glycosylation |
| GO:0006457 | protein folding |
| GO:0006091 | generation of precursor metabolites and energy |
| GO:0005975 | carbohydrate metabolic process |
| GO:0003016 | respiratory system process |
| GO:0003014 | renal system process |
| GO:0003013 | circulatory system process |
| GO:0003012 | muscle system process |
| GO:0002376 | immune system process |
| GO:1901135 | carbohydrate derivative metabolic process |
| GO:0071941 | nitrogen cycle metabolic process |
| GO:0061007 | hepaticobiliary system process |

#### A.5 Human-readable outputs

We provide here further detail on the final stage of PHILHARMONIC, using an LLM to summarize cluster functions into a human-readable output.

##### A.5.1 LLM Prompt

```
task_instruction = """
You are an expert biologist with a deep understanding of the Gene Ontology. Your job is to give
    short, intuitive, high-level names to clusters of proteins, given a set of GO terms
    associated with the proteins and their frequencies.
"""

confidence_score = """
In addition to the name, you should indicate how confident you are that your label is correct,
    and that it is representative of the function of the cluster. This score should be None, Low,
     Medium, or High. If you cannot find a connection between the functions in the cluster, give
    a name of Unknown with a confidence of None.
"""

format_instruction = """
Your response should start with only the name of the cluster on the first line. Then, provide a
    short paragraph including your explanation for why you gave the cluster that name. Finally,
    on a new line, provide your confidence score. You should not put blank lines between these
    sections.
"""

analytical_approach = """
You should try to be as specific as possible in your naming, to avoid overlapping names with
    other similar clusters. However, the names should still be short and human readable, ideally
    fewer than 10 words. You should consider the most common GO terms in the cluster, and try to
    find a common theme or function that ties them together. If you cannot find a common theme,
    you should give the cluster a name of Unknown.
"""

one_shot_example = """
For example, given the cluster description below:

Cluster of 14 [pdam_00002129-RA,pdam_00001718-RA,...] (hash 1332063120138743063)
```

```
Triangles: 27.0
Max Degree: 8
Top Terms:
    GO:0071502 - <cellular response to temperature stimulus> (11)
    GO:0019233 - <sensory perception of pain> (11)
    GO:0042493 - <response to drug> (10)
    GO:0007603 - <phototransduction, visible light> (10)
    GO:0004876 - <complement component C3a receptor activity> (9)

We would name this cluster Temperature, Pain, and Drug Response because there is a high representation for GO terms related to temperature, drug, and pain response.
"""

request = """
Please name the following cluster:
"""

LLM_SYSTEM_TEMPLATE = (
    task_instruction
    + confidence_score
    + format_instruction
    + analytical_approach
    + one_shot_example
    + request
)
```

##### A.5.2 Sample PHILHARMONIC JSON cluster.

The main output of PHILHARMONIC is a `.json` file containing a full specification of each cluster, including all members, proteins re-added by ReCIPE, the subgraph of edges in the cluster, GO term annotations, and all human-readable annotations.


```
...,
"208641124039621440": {
        "members": [
            "pdam_00013683-RA",
            "pdam_00006515-RA",
            "pdam_00000216-RA",
            "pdam_00009314-RA",
```

```
            "pdam_00024660-RA",
            "pdam_00021435-RA",
            "pdam_00000370-RA",
            "pdam_00023856-RA",
            "pdam_00022321-RA",
            "pdam_00016148-RA",
            "pdam_00006995-RA",
            "pdam_00019541-RA",
            "pdam_00014380-RA",
            "pdam_00000035-RA",
            "pdam_00003202-RA",
            "pdam_00007455-RA",
            "pdam_00017375-RA",
            "pdam_00006721-RA",
            "pdam_00003531-RA",
            "pdam_00022374-RA"
        ],
        "graph": [
            [
                "pdam_00022374-RA",
                "pdam_00023856-RA",
                0.5334205031394958
            ],
            [
                "pdam_00022374-RA",
                "pdam_00000035-RA",
                0.6888012290000916
            ],
            [
                "pdam_00007455-RA",
                "pdam_00023856-RA",
                0.6809149384498596
            ]
        ],
        "recipe": {
            "degree": {
                "0.75": [
                    "pdam_00021087-RA",
                    "pdam_00012633-RA",
```

```
                    "pdam_00011773-RA",
                    "pdam_00012527-RA",
                    "pdam_00003878-RA",
                    "pdam_00018049-RA",
                    "pdam_00018748-RA",
                    "pdam_00002992-RA",
                    "pdam_00021189-RA",
                    "pdam_00017594-RA",
                    "pdam_00013619-RA",
                    "pdam_00008678-RA",
                    "pdam_00006058-RA",
                    "pdam_00002321-RA",
                    "pdam_00015309-RA",
                    "pdam_00020434-RA",
                    "pdam_00008656-RA",
                    "pdam_00014328-RA",
                    "pdam_00011829-RA",
                    "pdam_00006524-RA"
                ]
            }
        },
        "GO_terms": {
            "GO:0030168": 18,
            "GO:0002032": 18,
            "GO:0022400": 18,
            .
            .
            .
            "GO:0008345": 2,
            "GO:0035269": 1,
            "GO:0046329": 1,
            "GO:0060049": 1
        },
        "human_readable": "Cluster Name: Cell Signaling and Regulation\nCluster of 20 proteins [
    pdam_00021685-RA, pdam_00003645-RA, pdam_00012637-RA, ...] (hash 2185119890364449780)\n20
    proteins re-added by ReCIPE (degree, 0.75)\nEdges: 6\nTriangles: 0\nMax Degree: 4\nTop Terms
    :\n\t\tGO:0030168 - <platelet activation> (19)\n\t\tGO:0002032 - <obsolete desensitization of
     G protein-coupled receptor signaling pathway by arrestin> (19)\n\t\tGO:0022400 - <regulation
     of opsin-mediated signaling pathway> (19)\n\t\tGO:0051586 - <positive regulation of dopamine
```

```
 uptake involved in synaptic transmission> (19)\n\t\tGO:0031635 - <adenylate cyclase-
    inhibiting opioid receptor signaling pathway> (19)\n\t\tGO:2000479 - <regulation of cAMP-
    dependent protein kinase activity> (19)\n\t\tGO:0040015 - <negative regulation of
    multicellular organism growth> (19)\n\t\tGO:0072224 - <metanephric glomerulus development>
    (19)\n\t\tGO:0070963 - <positive regulation of neutrophil mediated killing of gram-negative
    bacterium> (19)\n\t\tGO:0035025 - <positive regulation of Rho protein signal transduction>
    (19)\nLLM Explanation: This cluster consists of proteins that are highly associated with
    various signaling pathways and regulatory processes, particularly related to cell activation
    and communication. The predominance of GO terms related to platelet activation, GPCR
    signaling, dopamine uptake regulation, and various regulatory mechanisms suggests a strong
    focus on how cells interact and respond to stimuli, which includes growth regulation and
    immune function as well.\nLLM Confidence: High\n",
        "llm_name": "Cell Signaling and Regulation",
        "llm_explanation": "This cluster consists of proteins that are highly associated with
    various signaling pathways and regulatory processes, particularly related to cell activation
    and communication. The predominance of GO terms related to platelet activation, GPCR
    signaling, dopamine uptake regulation, and various regulatory mechanisms suggests a strong
    focus on how cells interact and respond to stimuli, which includes growth regulation and
    immune function as well.",
        "llm_confidence": "High"
    }...
```

##### A.5.3 Sample PHILHARMONIC human-readable output.

All clusters are presented to the end user in a flat text file. The `hash` allows the cluster to be identified in the accompanying `.json` file. These summaries allow a user to easily scan through clusters and find communities they are interested in investigating further.

```
Cluster Name: Pain, Drug Response, and Development
Cluster of 20 proteins [pdam_00013683-RA, pdam_00006515-RA, pdam_00000216-RA, ...] (hash
    208641124039621440)
20 proteins re-added by ReCIPE (degree, 0.75)
Edges: 3
Triangles: 0
Max Degree: 2
Top Terms:
        GO:0019233 - <sensory perception of pain> (20)
        GO:0048148 - <behavioral response to cocaine> (19)
        GO:0006468 - <protein phosphorylation> (19)
```

**Table A2: PHILHARMONIC predicted network statistics.** We run PHILHARMONIC on three species—the fruit fly *D. melanogaster*, the coral *P. damicornis*, its symbiont *C. goreaui*. Here, we show basic statistics of the predicted network for each species.

| | *P. damicornis* | *C. goreaui* | *D. melanogaster* |
|---|---|---|---|
| **Proteins** | 7,267 | 8,204 | 4,192 |
| **Edges** | 348,278 | 568,536 | 197,510 |
| **Med. Degree** | 37 | 44.5 | 45.5 |
| **Avg. Degree** | 95.852 | 138.560 | 94.232 |
| **Density** | 0.00656 | 0.00845 | 0.01124 |

```
        GO:0007507 - <heart development> (19)
        GO:0010759 - <positive regulation of macrophage chemotaxis> (19)
        GO:0001963 - <synaptic transmission, dopaminergic> (19)
        GO:0071380 - <cellular response to prostaglandin E stimulus> (19)
        GO:0071502 - <cellular response to temperature stimulus> (19)
        GO:0008542 - <visual learning> (19)
        GO:0007601 - <visual perception> (19)
LLM Explanation: This cluster is characterized by a strong representation of GO terms associated
    with sensory perception of pain, responses to drugs (specifically cocaine), and developmental
     processes, especially in the heart. The presence of terms related to sensory response and
    nervous system functions, alongside those concerning cellular processes and behavioral
    responses, suggests a common role in both response to stimuli and the development of certain
    physiological traits.
LLM Confidence: High
```

#### A.6 Supplemental material for the *D. melanogaster* analysis

We show detailed statistics of the fruit fly network in Table A2. In the main text, we perform a case study of a cluster in fly that demonstrates the additional value that PHILHARMONIC brings in addition to node-level functional annotation. Below, we show a full list of this cluster membership, both the gene name and the FlyBase identifier.

- 7227.FBpp0070550: AstA-R1
- 7227.FBpp0072053: G$\alpha$s
- 7227.FBpp0076643: G$\alpha$i
- 7227.FBpp0085065: DJ-1$\beta$
- 7227.FBpp0086741: DJ-1$\alpha$
- 7227.FBpp0087361: G$\alpha$o

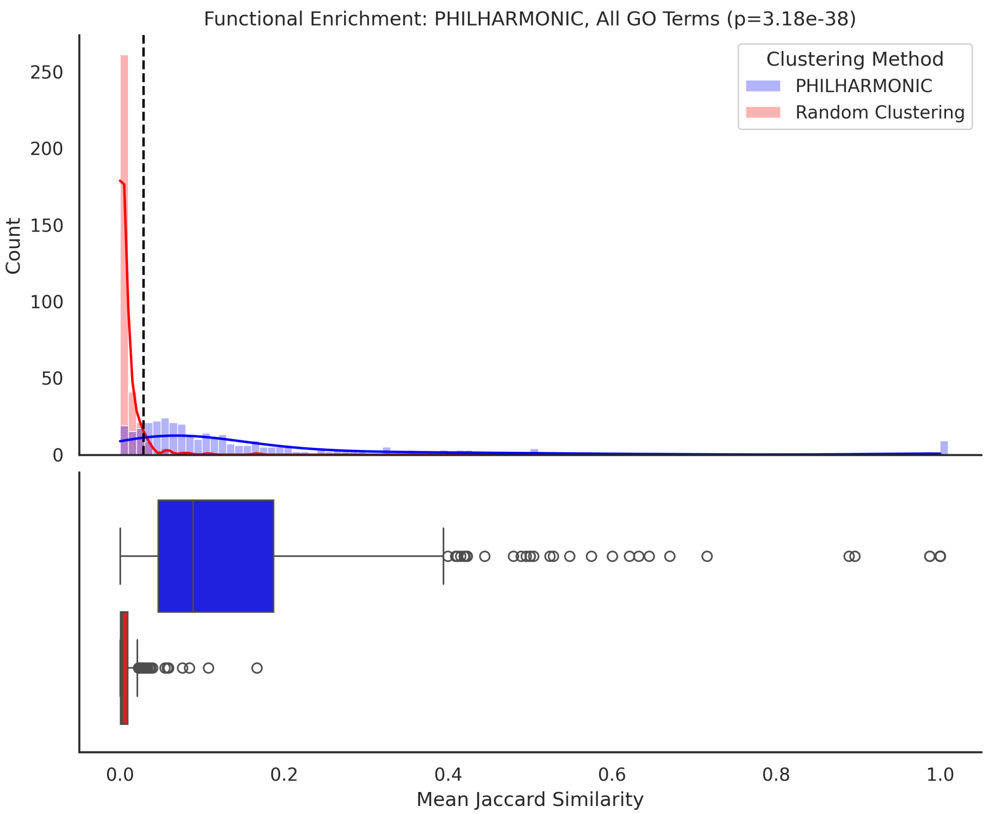


**Figure A2: PHILHARMONIC performs better with a higher-confidence network.** Compared to the results in Figure 2b, PHILHARMONIC clusters are significantly more coherent when the true STRING fly PPI network is used, rather than a fly network predicted using D-SCRIPT. We emphasize that this is not a possibility for many species, where a high-confidence PPI network is not available.

- 7227.FBpp0110121: G$\alpha$q
- 7227.FBpp0291583: G$\alpha$q
- 7227.FBpp0300610: Concertina

##### A.6.1 Effect of network noise on cluster functional coherence

While D-SCRIPT has been shown to achieve high accuracy in cross-species PPI prediction [17], it is likely that the predicted network contains noisy or missing edges. To isolate the effects of our downstream framework from the potential of noise in predicted edges, we substitute the network prediction step of PHILHARMONIC with the gold-standard *D. melanogaster* PPI network from STRING [7]. In Figure A2, we show the results of our clustring and functional coherence analysis on this network. As expected, clusters display stronger coherence when the underlying network is less noisy, underscoring the need for continued improvement of PPI prediction methods and the potential for additional performance gains of the PHILHARMONIC method.

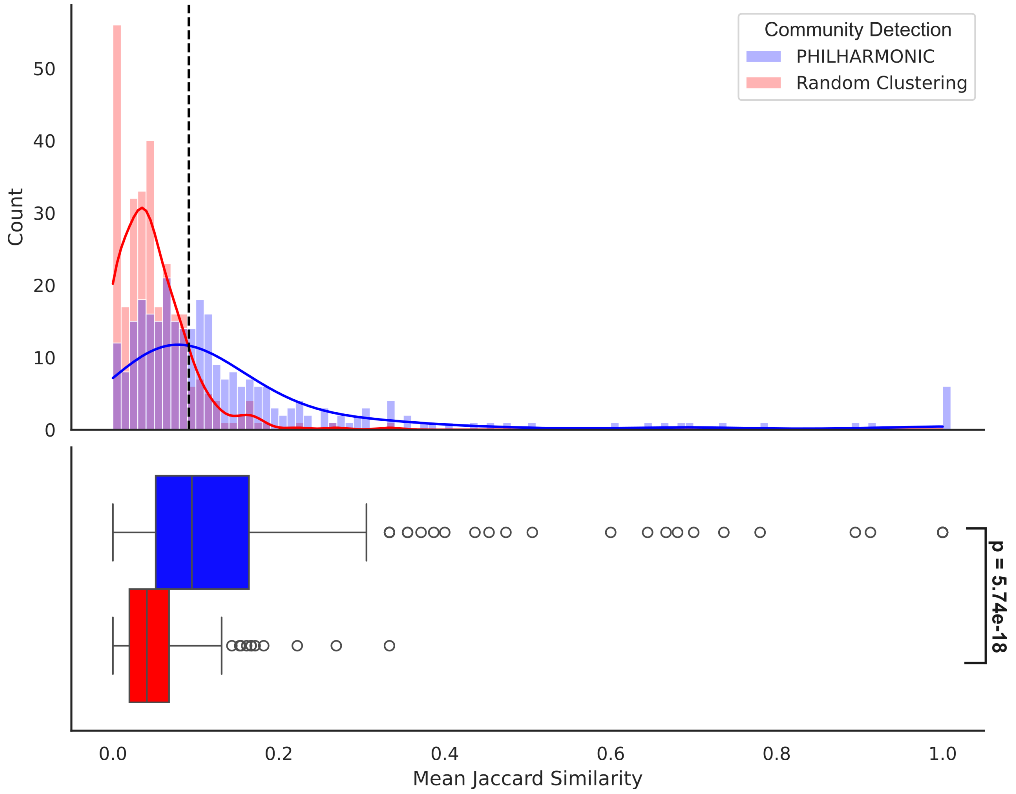


**Figure A3: PHILHARMONIC clusters group genes with shared FlyBase pathways.** Similar to Figure 2b, but using FlyBase pathways rather than gold-standard GO terms to determine protein relatedness. *D. melanogaster* proteins in a PHILHARMONIC cluster are signficantly more likely than a random background to be labeled with the same pathway in FlyBase.

##### A.6.2 Functional coherence using FlyBase pathway assignments

In addition to GO annotations, we also investigate a higher level of function. FlyBase reports gene groups such as "Proton-Coupled Amino Acid Transporters" or "Paired Homeobox Transcription Factors". We perform a similar analysis wherein we identify 853 gene groups annotated for at least one protein in our network. Then, we compute similarity within a cluster by the Jaccard similarity between sets of FlyBase gene groups for each protein, referred to here as "shared pathways." We show in Figure A3 that PHILHARMONIC clusters are significantly more likely to contain pairs of genes with shared biological pathways than random ($p = 5.74 \times 10^{-18}$, one-tailed independent t-test).

##### A.6.3 Gene expression analysis in *D. melanogaster*

We analyze co-expression of genes within PHILHARMONIC clusters using data from Schlamp et al. [28]. To ensure consistency across all species, we follow the data pre-processing and correlation procedures established in Connely et al. [44]; we compute gene co-expression scores by first computing a variance-stabilizing transform on expression values, then compute the bi-weighted mid-correlation statistic between genes (Figure 2c,d). Genes with significant

**Table A3: Use of other community detection methods for network clustering.** Only regularized spectral clustering (similar to a subroutine of our approach) and IPCA approach clustering results competitive with PHILHARMONIC. We show both methods for strictly non-overlapping clusters (first section) and methods which allow overlap (second section).

| Clustering Method | Number of Clusters | p-value |
|---|---|---|
| **RDS** | 468 | 2.668e-48 |
| Eigenvector [86] | 6 | 1.420e-01 |
| Infomap [87] | 53 | 9.908e-07 |
| Label propagation [88] | 27 | 1.590e-03 |
| Leiden [89] | 8 | 6.022e-02 |
| Louvain [90] | 9 | 6.342e-02 |
| Regularized spectral [91] | 440 | 7.578e-41 |
| Congo [92] | *Out of memory* | *>350 GB* |
| IPCA [93] | 2312 | 0.0 |
| LFM [94] | *Time out* | *>24 hours* |
| Multicom [95] | 6 | 2.990e-02 |
| Walkscan [96] | 1 | - |

missing data (0 counts in $> \frac{1}{2}$ of samples) were removed. As a baseline, we re-shuffle expression, preserving the distribution of expression values.

#### A.7 Supplemental material for the *P. damicornis* analysis

We show detailed statistics of all three networks, including numbers of nodes, edges, median and average degrees, and sparsity in Table A2.

##### A.7.1 Functional coherence analysis

In Figure 4c we compute the Jaccard similarity between pairs of proteins over the full set of GO terms. The Jaccard similarity is defined as

$$J(A, B) = \frac{A \cap B}{A \cup B} \tag{3}$$

where A, B are the sets of GO terms assigned to each individual protein. In Figure A4, we use only GO Slim terms to compute similarity between proteins. We find that even over that reduced set, PHILHARMONIC clusters are still significantly more functionally coherent than would be expected at random.

To study coherence of gene expression in *P. damicornis*, we use data from Connelly et al. [44]. We follow the same pre-processing procedure as for the fly analysis described above (Appendix A.6.3).

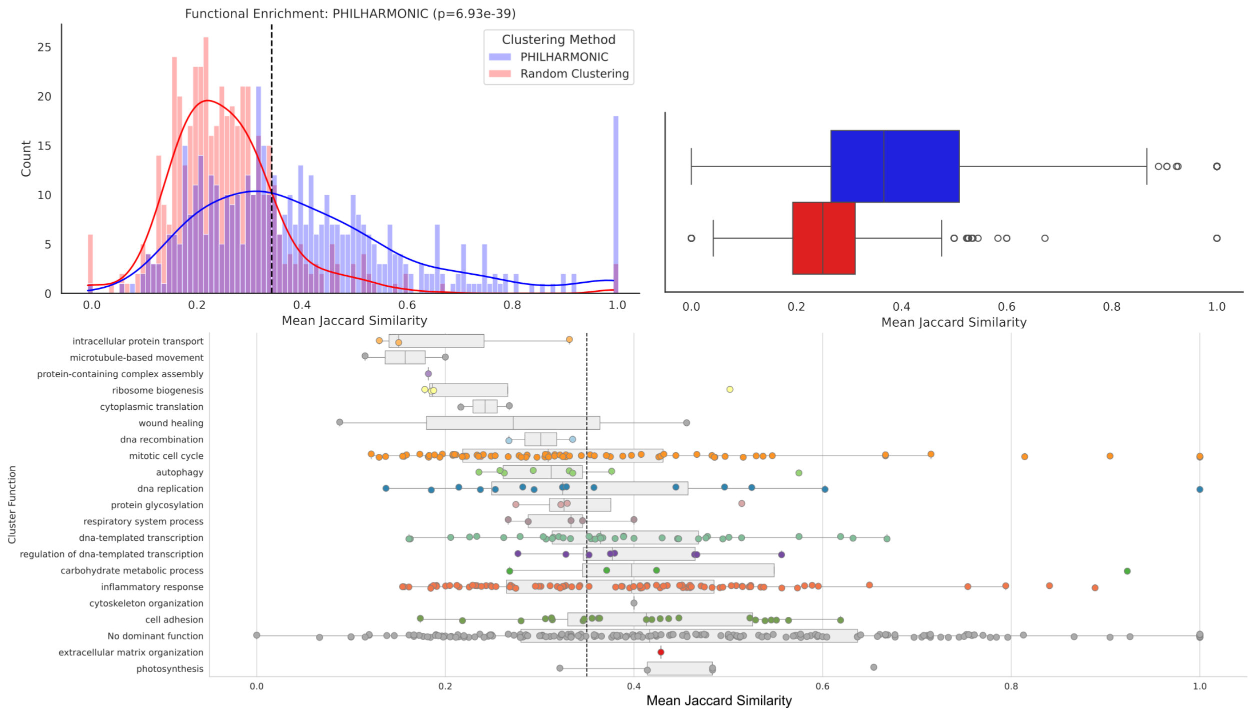


**Figure A4: *P. damicornis* functional coherence analysis using limited set of GO Slim terms.** We perform the same analysis as in Figure 4c using only the subset of GO terms from GO Slim. We find that PHILHARMONIC clusters are still significantly functionally enriched, although with a higher threshold to clearly separate non-random cluster coherences.

##### A.7.2 Comparison with other clustering methods

We previously showed that RDS yields clusters that are significantly functionally coherent. Here, we compare with several other state-of-the-art community detection methods, evaluating them by the same metric—the extent to which clusters detected by the given algorithm share more function that would be expected by random clustering. This analysis is performed on the *P. damicornis* network.

Using the CDlib Python package (version 0.4.0) [58], we evaluate several different algorithms for generating both overlapping and non-overlapping clusters. In Table A3, we show both the number of clusters detected by this method, and the $p$-value of the one-tailed t-test evaluating cluster coherence of computed vs. random clusters. Only regularized spectral clustering [91] and IPCA [93] have performance competitive with ours. In Figure A5, we show the distributions of cluster sizes and cluster coherences for each method. Only our method produces clusters with uniform size at a size which is easily amenable to biological discovery—both regularized spectral and IPCA result in many very large or small clusters that are difficult to interpret. We quantify the uniformity of cluster sizes by computing the entropy of the cluster size distribution (Table A4); RDS achieves the second-highest entropy indicating highly balanced cluster sizes. Only IPCA has a higher entropy, but it generates several very large clusters, and is computationally intractable on large networks [22].

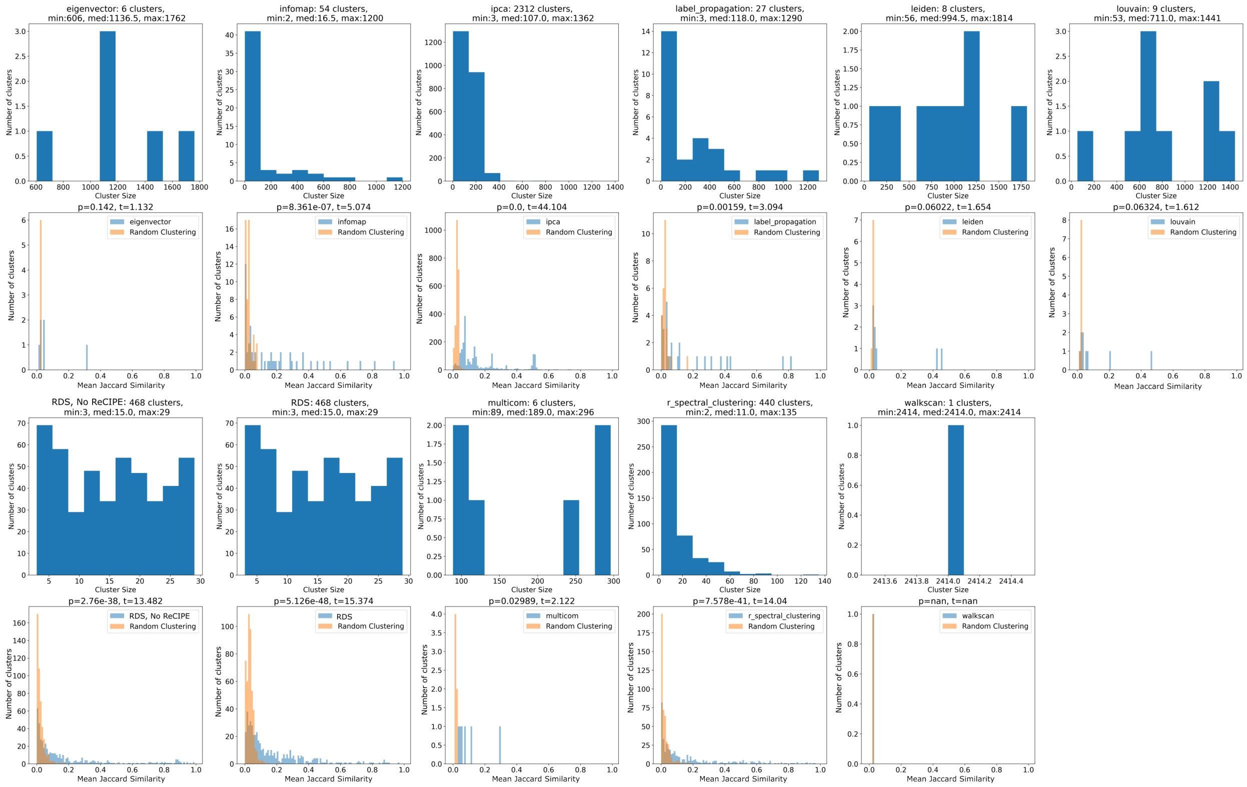


**Figure A5: Comparison of cluster sizes with other community detection methods.** Our RDS clustering approach yields both highly balanced and highly functionally coherent clusters. While IPCA and spectral clustering have competitive functional coherences, the distribution of cluster sizes makes it much more difficult to investigate the biological relevance of a given cluster.

**Table A4: RDS generates a highly balanced distribution of cluster sizes.** RDS cluster size entropy is the second highest behind only IPCA.

| Clustering Method | Entropy of Cluster Size Distribution |
|---|---|
| IPCA | 7.515237607193031 |
| **RDS** | 6.000553405179423 |
| Spectral Clustering | 5.729948575706442 |
| Infomap | 2.909253745840486 |
| Label Propagation | 2.649317997554160 |
| Louvain | 2.051001787760691 |
| Leiden | 1.878247804360152 |
| Eigenvector | 1.744732538032714 |
| MULTICOM | 1.672430619388146 |

##### A.7.3 Comparison with direct functional clustering

An alternative method for discovering functional communities within an organism would be to forgo the network step and directly cluster on functional annotations. To investigate this hypothesis, we develop an alternative clustering method that uses Jaccard similarity of assigned GO functional terms as a distance metric between proteins, and performs Hierarchical Spectral Clustering on this distance matrix. Because these clusters by definition have a high functional coherence, we evaluate the quality of clusters using the previously-defined co-expression metrics (median bi-weight mid correlation). We compare the function-derived clusters with PHILHARMONIC clusters, which we note do not take into account functional information at all, and simply rely on predicted network topology.

We find no significant difference in the distribution of co-expression for genes within function-derived clusters versus clusters from PHILHARMONIC ($p = 0.747$, Figure A6a). This suggests that PHILHARMONIC clusters are as likely on average to group co-expressed genes, than if one were to directly cluster based on function. However, PHILHARMONIC clusters have the added benefit of containing physically predicted PPIs; we quantify this by computing the number of edges in the subgraph of the full PPI network induced by each cluster, and by the number of connected components in the same subgraph. We find that clusters based solely on function tend to be highly disconnected, with very few edges in the induced subgraph (Figure A6b, left) and a large number of connected components (Figure A6b, right) compared to PHILHARMONIC clusters.

##### A.7.4 Singular value dominance of gene expression within clusters

In Figure 4d, we compute the coherence of a cluster by the pairwise co-expression of proteins within the cluster. Here, rather than looking at the average of several pairwise correlations, we instead subset the gene expression to include only the genes corresponding to proteins within a given cluster, and compute the singular values of this matrix. Then, we look at the dominance of the first singular value—where a high first singular value corresponds with strongly coordinated expression within the cluster. In Figure A7, we show that PHILHARMONIC clusters likewise have significantly more coordinated expression than random clusters.

##### A.7.5 Global mixing of GO slim function

In Figure 4a, we observe that some functions appear to be associated within “neighbhorhoods,” while others are dispersed around the network. To quantitatively assess the level of hierarchical organization within the cluster graph, we compute the node attribute assortativity over both the entire cluster graph and the edge weight-filtered cluster graph (shown in Figure 4a). Attribute assortativity measures the tendency of nodes to connect to other nodes with the same attribute ([-1, 1], higher is better) [97].

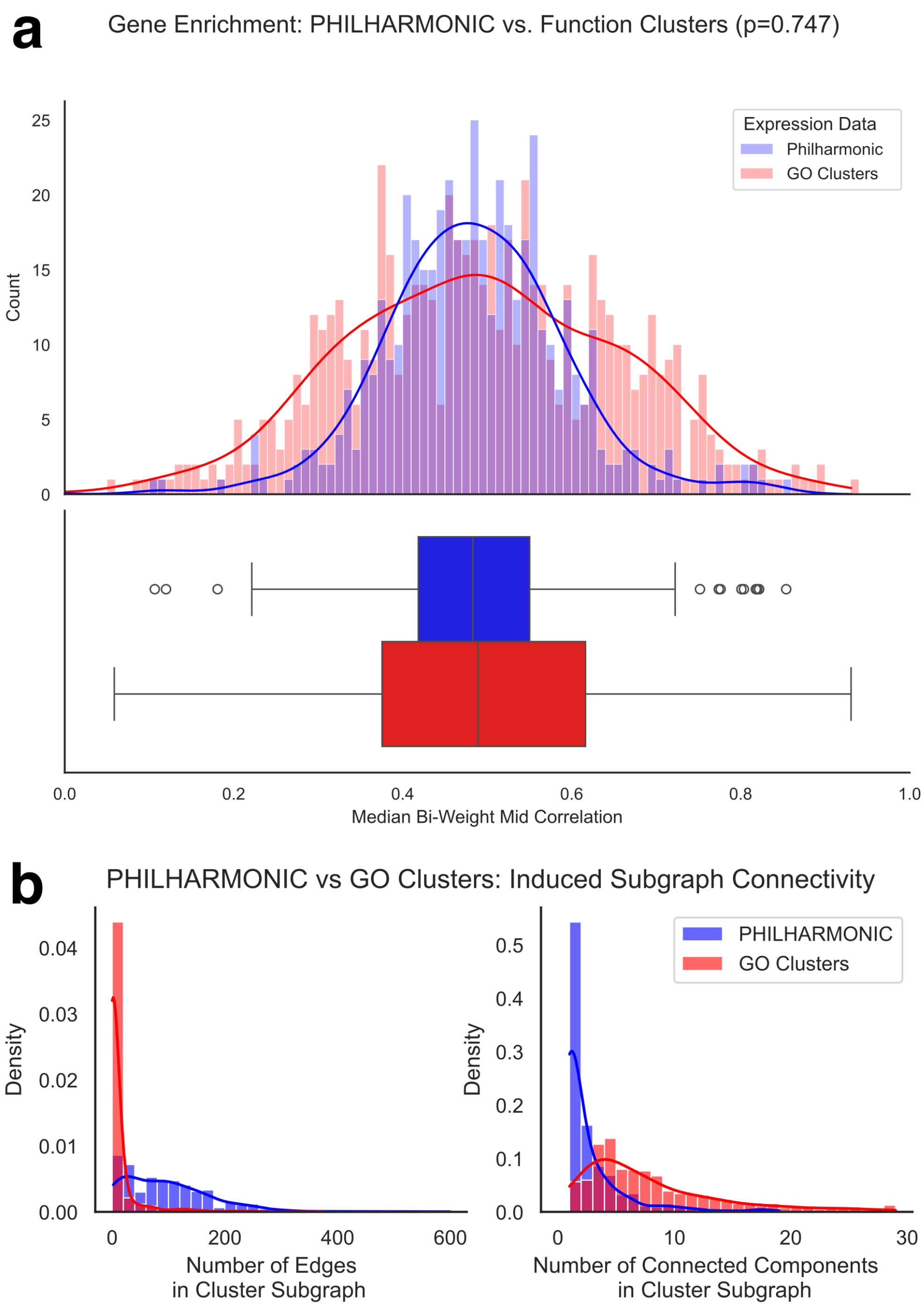


**Figure A6: Comparison of PHILHARMONIC with function-based clusters.** We create an alternate clustering by treating the Jaccard similarity between GO vectors as the distance between proteins, and performing hierarchical spectral clustering directly on this distance matrix. This yields a clustering with a similar size distribution. **(a)** These alternate clusters are no more likely than those from PHILHARMONIC to group co-expressed genes (independent samples T-test, p=0.747), using data from Connelly et al. **(b)** However, these alternate clusters are much more disconnected, with fewer PPIs between cluster members and substantially more connected components within each subgraph.

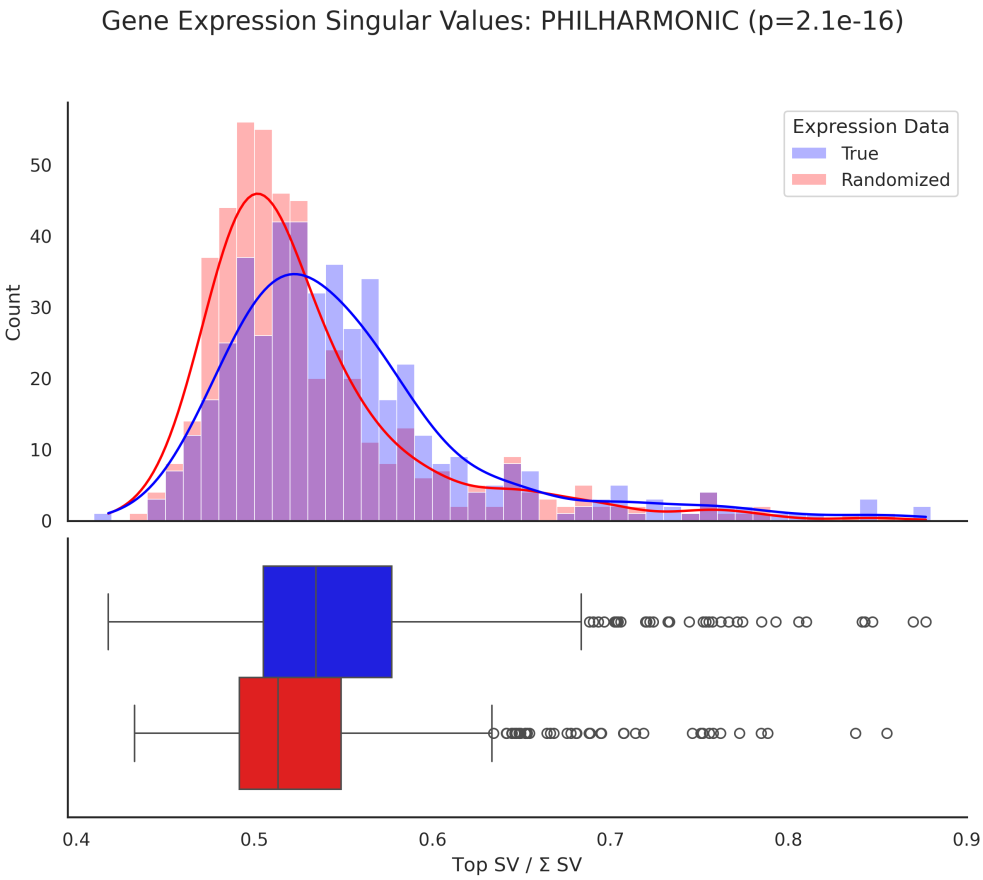


**Figure A7: Cluster coherence by singular value of gene expression** Similar to Figure 4d, but with co-expression calculated using the dominance of the first singular value.

The full graph cluster graph has an assortativity of 0.237 (0.296 weighted), while the filtered graph has an assortativity of 0.477 (0.397 weighted). Because the cluster graph is based on the number of crossing edges, the full graph is likely to have several very low-weight edges where a small number of PPIs occur across many clusters; we therefore think the filtered graph is a better representation of the true cluster connectedness. This assortativity supports the claim that clusters are in general well-connected to those with similar functions.

To evaluate the functional tightness of specific functions, we compute the conductance of that set [98], a measure of how well a set of labels is separated from the rest of the graph ([0, 1], lower is better, Table A5). We find that inflammatory response (0.097, 0.041 weighted), DNA-templated transcription (0.401, 0.262 weighted), and mitotic cell cycle (0.467, 0.404 weighted) are all well connected. In contrast, DNA replication (0.754, 0.794 weighted) and regulation of DNA-templated transcription (0.855, 0.880 weighted) are more distributed. We have updated the text to reflect this experiment and these new insights, clarifying which functions are mixed and which are coherent.

**Table A5: Conductance of GO term sets in the network.** Lower values indicate tighter functional connectivity.

| Function | GO Term | Conductance | Weighted Conductance |
|---|---|---|---|
| Inflammatory response | GO:0006954 | 0.097 | 0.041 |
| DNA-templated transcription | GO:0006351 | 0.401 | 0.262 |
| Mitotic cell cycle | GO:0000278 | 0.467 | 0.404 |
| DNA replication | GO:0006260 | 0.754 | 0.794 |
| Regulation of DNA-templated transcription | GO:0006355 | 0.855 | 0.880 |

###### A.7.6 Thermal response cluster

The majority of proteins in this cluster are predicted to be GPCRs, which are predicted to bind to various subsets and likely modulate the functions of the diverse ion channels in response to different stimuli. These proteins include `pdam_00021189`, with a sequence signature somewhat similar to the alpha-2Da adrenergic receptor detected in the zebrafish brain [99], and `pdam_00001720`, predicted by sequence homology to be an allatostatin-A receptor. The neuropeptide allatostatin-A (AstA) and its cognate receptors (AstARs) are involved in the modulation of feeding behavior in mosquitos. Two of the other proteins `pdam_00006261` (most likely an orexin receptor) and `pdam_00013140` (neuropeptide FF receptor 2-like) are both similar to receptor proteins that regulate the feeding process in fish [100]. `pdam_00023837`, `pdam_00016481` and `pdam_00016115` round out the collection of GPCRs. Finally, there are three proteins in the cluster which are not either ion channels or GPCRs: `pdam_00019706`, predicted to be a fibroblast growth factor, `pdam_00004445`, predicted to be a zinc transporter, and `pdam_00017094`, a poorly characterized protein that is most similar structurally to Nodulation protein Z or Alpha-(1,6)-fucosyltransferase by a FoldSeek search.

Below, we show a full list of cluster membership and their best homology match from a BLAST search (description and gene symbol). We note that these are not confirmed functions or genes, but rather the closest human sequence match. These descriptions and gene symbols are provided as additional context and are not part of the PHILHARMONIC output. One protein is **poorly characterized** by sequence homology, and we instead provide the best structural hit using FoldSeek in **bold.** They were not used in the construction of these clusters and were manually identified.

- pdam_00001720: allatostatin-A receptor-like (AstA-R1)
- pdam_00004445: zinc transporter 8-like (ZNT8)
- pdam_00005806: potassium/sodium hyperpolarization-activated cyclic nucleotide-gated channel 2-like (HCN2)
- pdam_00006261: orexin receptor type 1-like (OX1)
- pdam_00008678: cyclic nucleotide-gated channel rod photoreceptor subunit alpha-like (CNGA1)
- pdam_00010576: TWiK family of potassium channels protein 7-like isoform X1 (TWK7)
- pdam_00013140: neuropeptide FF receptor 2-like (NPFF2)
- pdam_00016115: beta-2 adrenergic receptor-like (ADRB2)
- pdam_00016481: beta-1 adrenergic receptor-like (ADRB1)
- pdam_00017094: **Nodulation protein Z (UniProt: Q45271)**
- pdam_00019465: potassium voltage-gated channel subfamily A member 7-like (KCNA7)
- pdam_00019542: potassium voltage-gated channel protein Shal-like isoform X1 (KCND2)
- pdam_00019706: fibroblast growth factor 2-like isoform X2 (FGF2)
- pdam_00021189: alpha-2Da adrenergic receptor-like (ADRA2A)
- pdam_00023837: histamine H2 receptor-like, partial (HRH2)

Finally, we show the PHILHARMONIC-produced human-readable output for this cluster:

```
Cluster Name: Temperature and Pain Regulation Cluster
Cluster of 15 proteins [pdam_00008678-RA, pdam_00021189-RA, pdam_00005806-RA, ...] (hash
    1495076087230339862)
```

```
0 proteins re-added by ReCIPE (degree, 0.75)
Edges: 31
Triangles: 8
Max Degree: 8
Top Terms:
        GO:0071502 - <cellular response to temperature stimulus> (12)
        GO:0019233 - <sensory perception of pain> (12)
        GO:0002024 - <diet induced thermogenesis> (12)
        GO:0042391 - <regulation of membrane potential> (9)
        GO:0043547 - <positive regulation of GTPase activity> (8)
        GO:0070374 - <positive regulation of ERK1 and ERK2 cascade> (8)
        GO:0030168 - <platelet activation> (7)
        GO:0002032 - <obsolete desensitization of G protein-coupled receptor signaling pathway by arrestin> (7)
        GO:0022400 - <regulation of opsin-mediated signaling pathway> (7)
        GO:0051586 - <positive regulation of dopamine uptake involved in synaptic transmission> (7)
LLM Explanation: This cluster is characterized by a strong presence of GO terms related to temperature response and sensory perception of pain, highlighting its connection to thermoregulation and nociception. Additionally, there are terms associated with the regulation of membrane potentials and various signaling pathways, which suggest a key role in cellular signaling and response mechanisms that may be triggered by changes in temperature and pain stimuli. Therefore, I have named this cluster based on its emphasis on temperature and pain-related functions.
LLM Confidence: High
```

##### A.7.7 Environmental stimuli cluster

Below, we show a full list of cluster membership and their best homology match from a manual BLAST search and analysis. We note that these are not confirmed functions or genes, but rather the closest human sequence match. Three proteins are **poorly characterized** by sequence homology, and we instead provide the best structural hit using FoldSeek in **bold.**

- pdam_00001381: potassium voltage-gated channel subfamily D member 3-like (KCND3)
- pdam_00001388: gamma-aminobutyric acid receptor alpha-like (GABRA1)
- pdam_00001389: gamma-aminobutyric acid receptor subunit alpha-2-like (GABRA2)

- pdam_00001924: glycine receptor subunit alphaZ1-like (GLRA1)
- pdam_00003290: transient receptor potential cation channel subfamily V member 6-like (TRPV6)
- pdam_00003714: histone deacetylase 1-like (HDAC1)
- pdam_00006109: **Follistatin-related protein 4 (UniProt: Q6MZW2-2)**
- pdam_00006156: calcium-independent phospholipase A2-gamma-like ( PNPLA8)
- pdam_00006320: potassium voltage-gated channel subfamily A member 2-like (KCNA2)
- pdam_00006972: neuronal acetylcholine receptor subunit alpha-10-like (CHRNA10)
- pdam_00006973: neuronal acetylcholine receptor subunit alpha-10-like isoform X1 (CHRNA10)
- pdam_00008620: FGFR1 oncogene partner 2 homolog (FGFR1OP2)
- pdam_00011550: **Pikachurin (Uniprot: Q4VBE4-2)**
- pdam_00011780: lipid droplet-associated hydrolase-like ( LDAH)
- pdam_00012376: gamma-aminobutyric acid receptor subunit beta-2-like (GABRB2)
- pdam_00012507: neuronal acetylcholine receptor subunit alpha-5-like (CHRNA5)
- pdam_00015999: neuronal acetylcholine receptor subunit alpha-7-like isoform X2 (CHRNA7)
- pdam_00018139: **Spondin (UniProt: Q9HCB6)**
- pdam_00021982: phosphatidylinositol phosphatase PTPRQ-like (PTPRQ)
- pdam_00024051: gamma-aminobutyric acid receptor alpha-like (GABRA1)

We also show the PHILHARMONIC-produced human-readable output for this cluster:

```
Cluster Name: Neurophysiological Responses to Environmental Stimuli
Cluster of 12 proteins [pdam_00001381-RA, pdam_00006156-RA, pdam_00012376-RA, ...] (hash
    803905878821692717)
8 proteins re-added by ReCIPE (degree, 0.75)
Edges: 8
Triangles: 0
Max Degree: 3
Top Terms:
```

```
        GO:0001964 - <startle response> (9)
        GO:0098703 - <calcium ion import across plasma membrane> (9)
        GO:0045188 - <regulation of circadian sleep/wake cycle, non-REM sleep> (9)
        GO:0071805 - <potassium ion transmembrane transport> (7)
        GO:0071361 - <cellular response to ethanol> (7)
        GO:0001988 - <positive regulation of heart rate involved in baroreceptor response to decreased systemic arterial blood pressure> (7)
        GO:0017085 - <response to insecticide> (7)
        GO:0035095 - <behavioral response to nicotine> (7)
        GO:0060013 - <righting reflex> (7)
        GO:0021771 - <lateral geniculate nucleus development> (7)
LLM Explanation: This cluster includes a variety of GO terms that relate to responses in the nervous system and actions involving ion transport, such as the startle response and behavioral responses to substances like ethanol and nicotine. Additionally, the inclusion of terms concerning ion transmembrane transport suggests a role in neural excitability, which is critical for responding to environmental stimuli. The overarching theme indicates a connection to neurophysiological responses, particularly those triggered by changes in the environment.
LLM Confidence: High
```

##### A.7.8 Temperature and pain regulation cluster

Here, we report another case where PHILHARMONIC provides additional functional information on top of sequence or structural homology. We show this cluster highlighted in Figure A8a, where it acts as a connecting point between the local neighborhood of immune response clusters and the rest of the graph. We show each individual protein in this cluster in Figure A8b. 14 of these proteins are annotated for GO:0071502, a response to temperature stimulus. Of the 16 proteins in the cluster, we were only able to find gene expression information for two (pdam_00001658, pdam_00002742).

Below, we show a full list of cluster membership and their best homology match from a manual BLAST search and analysis against human proteins. We note that these are not confirmed functions or genes, but rather the closest human sequence match. Two proteins are **poorly characterized** by sequence homology, and we instead provide the best structural hit using FoldSeek in **bold.** Even by structural homology, these proteins are poorly characterized. pdam_00002663 has several Foldseek hits for Adensoine Receptor A1, but none are a lower e-value than $3.04e-2$, and the predicted AlphaFold structure used for Foldseek search has an average pLDDT of 61.97. pdam_00003116 has several Foldseek hits for Nephronectin, but none lower than e-value of $1.49e-3$, and the structure has an average pLDDT of 68.81.

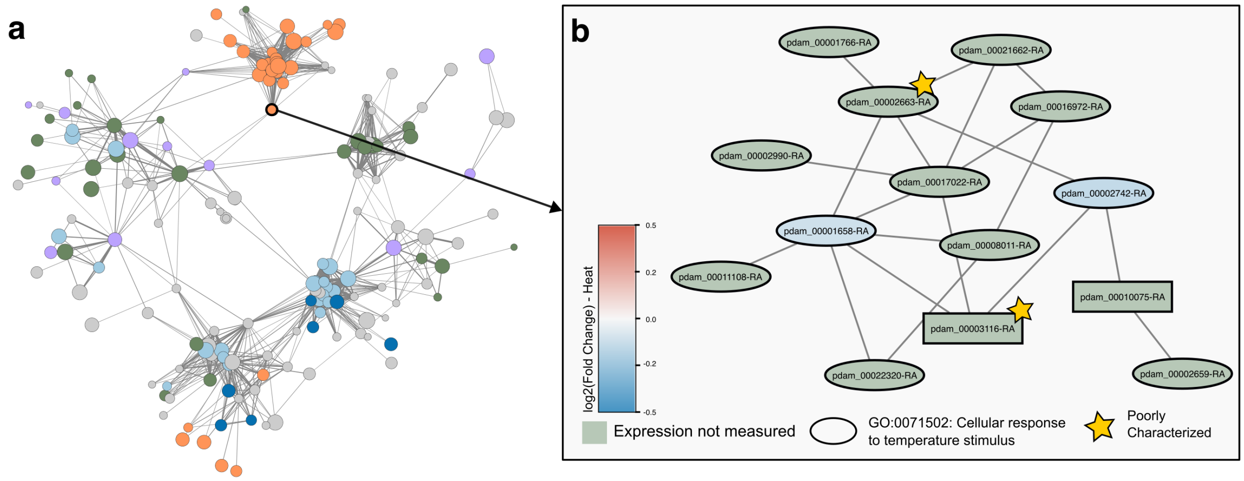


**Figure A8: Temperature and pain regulation in *P. damicornis*.** We show here a cluster of 16 proteins annotated for temperature and pain regulation. **(a)** This cluster is fairly central in the broader *P. damicornis* network, connecting the neighborhood of immune response clusters to the broader network. **(b)** This cluster contains 14 proteins annotated for GO:0071502 (cellular response to temperature stimulus), and two proteins that show moderately decreased expression in response to heat stress. Two proteins (stars) are poorly characterized by both sequence and structural homology.

- pdam_00001658: Adenosine receptor A2a (ADORA2A)
- pdam_00001766: Potassium voltage-gated channel subfamily A member 1 (KCNA1)
- pdam_00002659: Melatonin receptor type 1A (MTNR1A)
- pdam_00002663: **Adenosine Receptor A1 (UniProt: P28190)**
- pdam_00002742: D(1A) dopamine receptor (DRD1)
- pdam_00002990: Visual pigment-like receptor peropsin (RRH)
- pdam_00003116: **Nephronectin (UniProt: Q91V88)**
- pdam_00008011: Visual pigment-like receptor peropsin (RRH)
- pdam_00010075: ATP-binding cassette (ABCC4)
- pdam_00011108: Melatonin receptor type 1A (MTNR1A)
- pdam_00016972: Melatonin receptor type 1A (MTNR1A)
- pdam_00017022: Orexin receptor type 2 (HCRTR2)
- pdam_00021662: Potassium channel subfamily K member 5 (KCNK5)
- pdam_00022320: Melanopsin isoform 1 (OPN4)

We also show the PHILHARMONIC-produced human-readable output for this cluster:

```
Cluster Name: Temperature and Pain Regulation
Cluster of 16 proteins [pdam_00019213-RA, pdam_00022320-RA, pdam_00010075-RA, ...] (hash 1337569685223779389)
0 proteins re-added by ReCIPE (degree, 0.75)
Edges: 20
Triangles: 5
Max Degree: 6
Top Terms:
        GO:0071502 - <cellular response to temperature stimulus> (14)
        GO:0019233 - <sensory perception of pain> (14)
        GO:0002024 - <diet induced thermogenesis> (14)
        GO:0030168 - <platelet activation> (11)
        GO:0002032 - <obsolete desensitization of G protein-coupled receptor signaling pathway by arrestin> (11)
        GO:0022400 - <regulation of opsin-mediated signaling pathway> (11)
        GO:0051586 - <positive regulation of dopamine uptake involved in synaptic transmission> (11)
        GO:0031635 - <adenylate cyclase-inhibiting opioid receptor signaling pathway> (11)
        GO:2000479 - <regulation of cAMP-dependent protein kinase activity> (11)
        GO:0040015 - <negative regulation of multicellular organism growth> (11)
LLM Explanation: This cluster includes a significant representation of terms associated with cellular responses to temperature and sensory perception of pain, indicating a potential role in mechanisms that deal with stressors and pain perception. Additionally, the presence of terms related to platelet activation and signaling pathways suggests involvement in regulatory processes at a cellular level, possibly linking these stress responses to signaling in the nervous system and vascular health.
LLM Confidence: High
```

#### A.8 Supplemental material for the *C. goreaui* network

We show the symbiont cluster graph in Figure A9c, along with node degree and cluster size distributions (Figure A9a,b). We show detailed statistics of the *C. goreaui* network in Table A2. Below, we replicate the functional coherence analysis and highlight two clusters related to cellular oxidant de-toxification.

There are two clusters (Cluster of 11 and Cluster of 19) that contain many putative stress response genes. In Figure A9 we show the PHILHARMONIC human-readable output (A9d,A9f) and cluster graphs (A9e,A9g). The cluster of 11 (A9e, orange) includes 3 probable glutathione S-transferase (GST) proteins (SymbC1.scaffold1814.1,

SymbC1.scaffold218.8, SymbC1.scaffold7418.2) as well as 5 likely glutaredoxin proteins (SymbC1.scaffold40305.1, SymbC1.scaffold8553.1, SymbC1.scaffold241.40, SymbC1.scaffold5815.2, SymbC1.scaffold17803.1). The cluster of 19 (A9g, yellow) includes 7 probable GST proteins (SymbC1.scaffold3459.6, SymbC1.scaffold6731.6, SymbC1.scaffold20.307, SymbC1.scaffold2148.2, SymbC1.scaffold10891.1, SymbC1.scaffold4197.2, SymbC1.scaffold497.1), and 3 possible additional GST proteins (SymbC1.scaffold2726.4, SymbC1.scaffold80.118, SymbC1.scaffold19.140), as well as 2 probable glutaredoxin proteins (SymbC1.scaffold3871.1, SymbC1.scaffold685.22). GST and glutaredoxin proteins are known to be part of antioxidant response pathways to oxidative stress [101, 102], with glutaredoxins hypothesized to play a particularly important role in response to photo-oxidative stress in photosynthesizing plants [103]. In addition to the GST and glutaredoxim proteins, the cluster of 11 also contains three cytochrome proteins (SymbC1.scaffold3718.5, SymbC1.scaffold5013.1, SymbC1.scaffold994.10),one of which is a likely cytochrome-450 protein, where this family of proteins is generally preserved from algae to higher plants and can oxidize endogenous substrates in various biosynthetic pathways as well as xenobiotic substrates, in particular herbicides [104]. Other proteins in the cluster of 19 (yellow) include proteins with remote homology to the YghU protein (SymbC1.scaffold3998.31, SymbC1.scaffold4.1591, SymbC1.scaffold688.7), one similar to the related yfcG protein, and a methionine sulfide reductase protein (SymbC1.scaffold21.80) These proteins all have been identified as modulating the response to oxidative stress, where YghU and yfcG appear to be a novel form of GST proteins [105], and methionine sulfide reductase proteins repair oxidized proteins and are protective against damage caused by oxidative stress [106]. We hypothesize that these two clusters are important for reaction to oxidative stress in the symbiont.

We perform the same analysis of cluster functional coherence using predicted GO terms in the symbiont *C. goreaui*. We likewise find significant functional coherence in PHILHARMONIC clusters ($p = 2.74 \times 10^{-44}$). We show the results of this analysis using all GO terms in Figure A10; we find similar results using GO Slim terms.

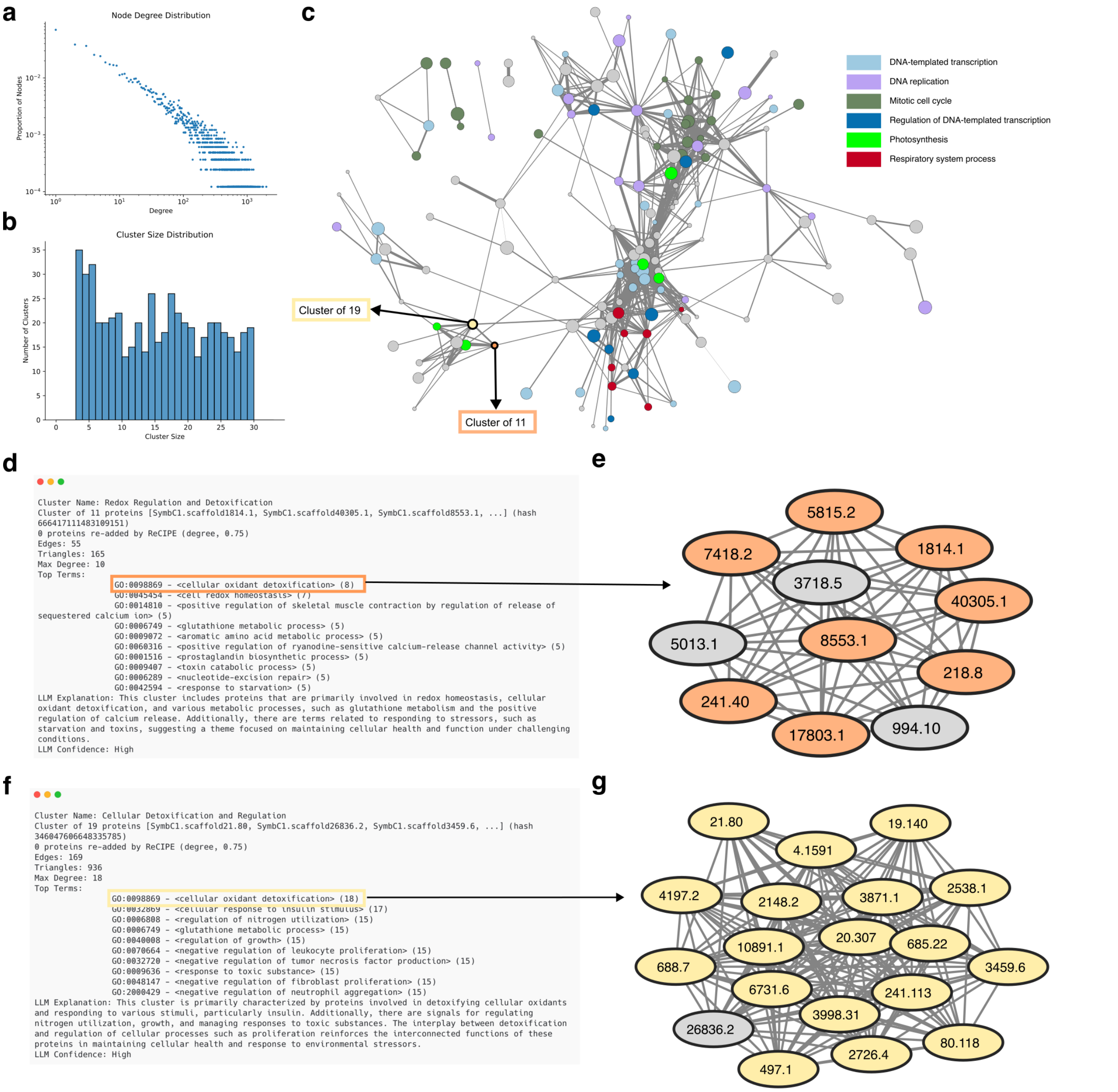


**Figure A9: Dissecting the functional network of the symbiont *C. goreaui*.** **(a)** Node degree distribution of predicted network. **(b)** Size distribution of PHILHARMONIC clusters. **(c)** Cluster graph, shown with $t = 50$ connecting edges. **(d)** Human-readable cluster description for the cluster of 11 described in Appendix A.8. For readability, we have dropped the `SymbC1.scaffold` prefix from node labels. **(e)** Graph for this cluster, colored by annotation for cellular oxidant detoxification (GO:0098869). **(f)** Human-readable cluster description for the cluster of 19 described in Appendix A.8. **(g)** Graph for this cluster, colored by annotation for cellular oxidant detoxification (GO:0098869).

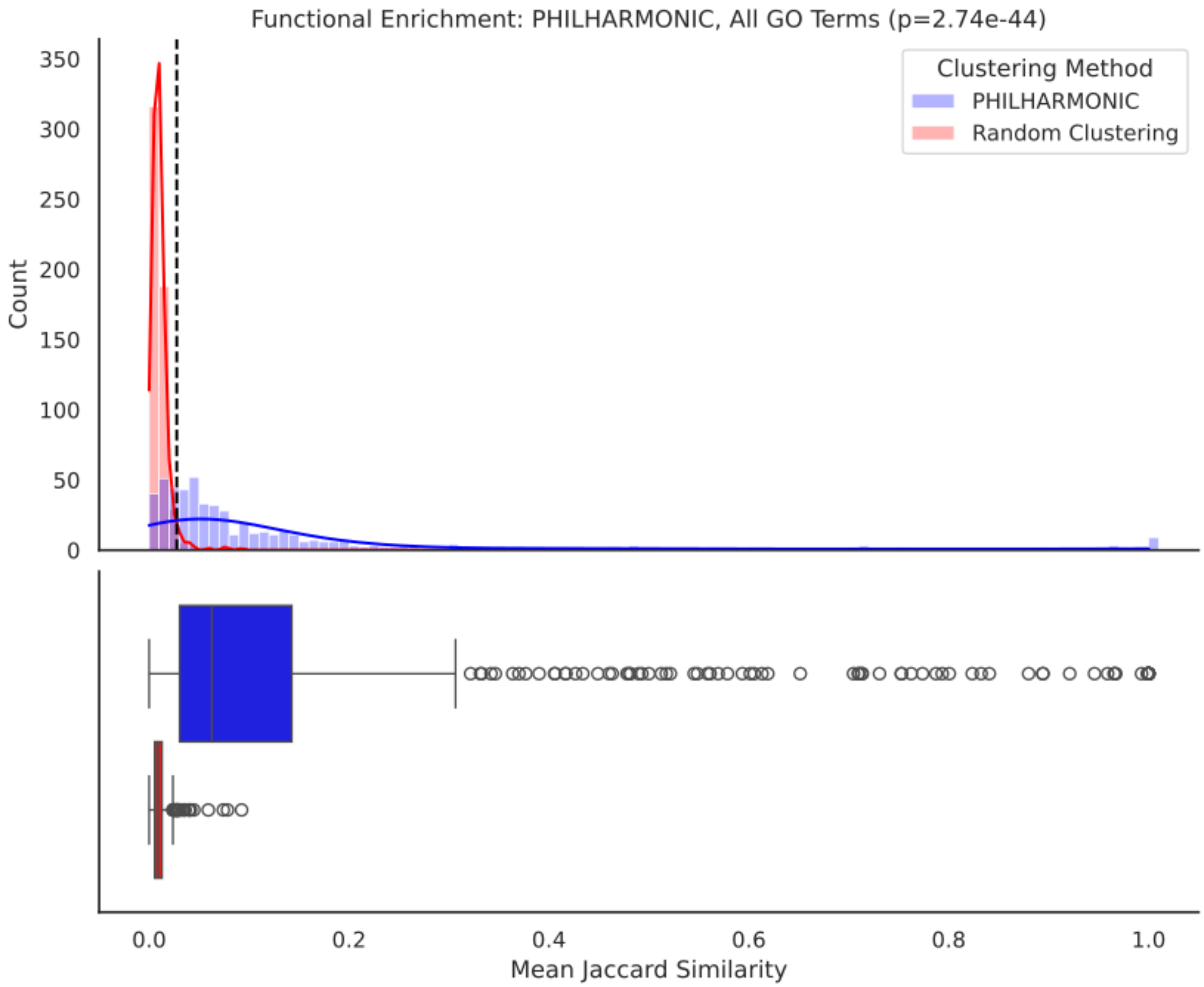


**Figure A10: Functional coherence analysis using all GO terms in *C. goreaui*.**

#### A.9 Pseudocode for the ReCIPE Algorithm

```
def reconnect(clusters, linear_ratio=0.1, cthresh=0.75, max_proteins=20, metric="min degree"):
    all_added_proteins = {}
    for cluster in clusters:
        added_proteins = {}
        potential:= proteins not in cluster
        sort potential proteins by metric

        num_proteins:= proteins in cluster

        num_components:= connected components in cluster
        percent_connectivity = 1 - (num_components - 1) / (num_proteins - 1)

        while (len(potential) > 0) and (percent_connectivity < cthresh):
            connection_minimum = linear_ratio * num_components

            for prot in potential:
                prot_degree:= number of components prot connects

                if prot_degree >= connection_minimum
                    add prot to added_proteins
                    recompute num_components
                    recompute percent_connectivity
                    remove prot from potential

                if len(added_proteins) > max_proteins:
                    break

        add (cluster, added_proteins) to all_added_proteins

    return all_added_proteins
```

#### A.10 RDS and ReCIPE validation

##### A.10.1 Evaluating robustness to hyperparameter settings in coral

We perform an in-depth evaluation of the quality of PHILHARMONIC clusters with a wide variety of different parameter settings both for clustering and for re-connection with ReCIPE. Specifically, we evaluate combinations of the following parameters over the *P. damicornis* network:

- Initial number of clusters: [50, 100, 500]
- Cluster divisor (at each iteration): [5, 10, 20]
- Minimum cluster size: [3, 10]
- ReCIPE linear ratio: [0.1, 0.25]
- ReCIPE cthresh: [0.25, 0.5, 0.75]
- ReCIPE maximum proteins re-added: [10, 20, 50]

We provide the full set results in the attached Supplementary Sheet S1. Broadly, we find that PHILHARMONIC yields functionally coherent clusters across many parameter settings, with the strongest performance coming when $lr = 0.1, cthresh = 0.75$. We do identify a failure mode for PHILHARMONIC—specifically, if the cluster division at each step is too small (5), while at the same time the minimum cluster size is too large (10), we end up creating many small clusters in the end, which are then filtered out resulting in no clusters being returned. Provided that a low enough minimum size is set (we recommend 3) or a large split at each step ($\geq$ 10), this outcome can be avoided. In future work, we will explore methods for automatically detecting and correcting this failure mode.

##### A.10.2 Evaluating robustness in yeast

While we showed that RDS and ReCIPE are broadly robust to parameter choice in coral, we next sought to evaluate whether these parameter choices were stable in other species. To do so, we use the *S. cerevisiae* (Baker's yeast) protein–protein interaction (PPI) network from STRING v11.5 [107], filtered to include only experimentally validated interactions with a combined confidence score $\geq$ 500. The resulting network comprises 2,394 proteins and 17,822 undirected edges. To evaluate the impact of network sparsity on RDS parameters, we also considered randomly down-sampled networks of size 10%, 25%, 50%, 75% and 90% through subsampling. Each threshold was subsampled 10 times. We report the Jaccard similarity and F1 score of clusters based on true GO functions in Figure A11. Here too, we find broadly consistent performance, with the best clustering achieved at $lr = 0.1, cthresh = 0.75$.

##### A.10.3 ReCIPE improves coherence on DREAM networks

As true information about functional clustering is sparse outside of human data, we additionally evaluate RDS and ReCIPE in three human PPI networks with known functions sourced from the Dialogue on Reverse Engineering

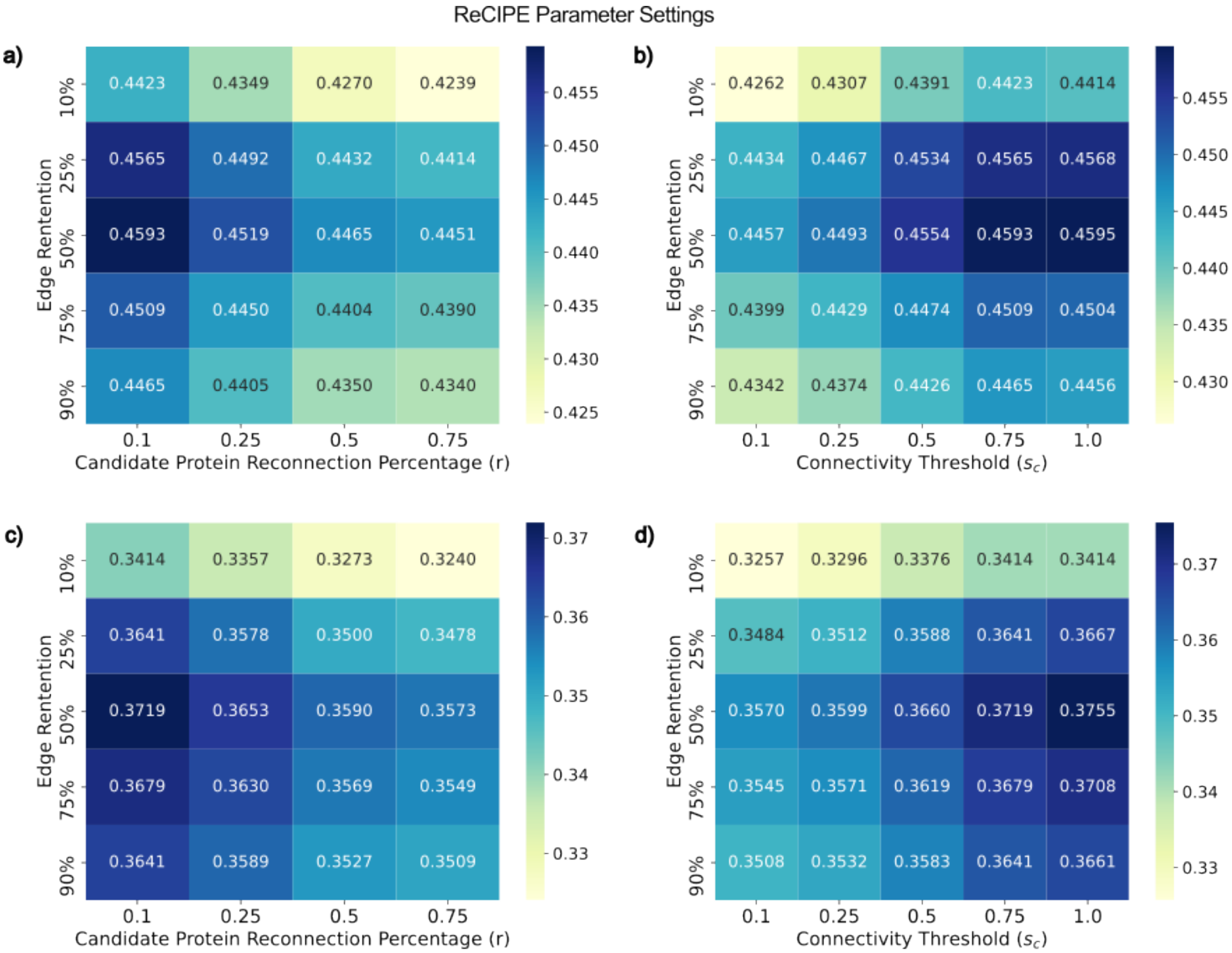


**Figure A11: Parameter optimization of ReCIPE for subsampled STRING network analysis.** Each panel displays the scores at various edge retention levels for subsampled STRING networks, averaged over 10 samplings. Panels a) and b) show F1 scores, as described in the Methods section, while panels c) and d) display Jaccard similarity. Panels a) and c) illustrate the effect of varying the candidate protein reconnection percentage ($r$) with the connectivity threshold fixed at 100% to maximize subgraph connectivity. Notably, $r = 0.1$ yields the highest F1 and Jaccard scores across all edge retention thresholds. Panels b) and d) demonstrate the impact of varying the connectivity threshold ($s_c$). For these data, $r$ was fixed at 0.1. The improved F1 and Jaccard scores achieved by increasing $s_c$ demonstrates how re-adding proteins that may have been assigned to different clusters in the spectral clustering step leads to improved functional metrics. Given the similarity of F1 and Jaccard scores between $s_c = 0.75$ and $s_c = 1.0$, we fixed $s_c = 0.75$ to minimize the number of re-added proteins.

Assessment and Methods (DREAM) challenge [1]. DREAM 1 is a PPI network representing protein interactions and functional association, which is derived from the STRING database [7]; the network has 17,388 proteins and 1,973,788 edges. DREAM 2 is a PPI network that only reflects physical protein networks, which is derived from the InWeb database [108]; the network has 12,325 proteins and 397,254 edges. DREAM 3 is a signaling network, which is curated from 27 sources from OmniPath [109], and contains 5,009 proteins and 18,270 edges. We show statistics for all three of these networks in Table A6.

**Table A6: Statistics of DREAM networks.**

| | **DREAM 1** | **DREAM 2** | **DREAM 3** |
|---|---|---|---|
| **Nodes** | 17397 | 12420 | 5009 |
| **Edges** | 2232405 | 397309 | 18424 |
| **Density** | 0.0150 | 0.00515 | 0.00134 |
| **Avg. clustering coeffecient** | 0.325 | 0.291 | 0.134 |
| **Avg. degree** | 257 | 64 | 7 |

In Figure A12, we show the function prediction results for all three networks computed by two methods. For both methods, we use the FUNC-E [110] package to compute a set of enriched GO terms for a cluster. We assign a cluster all enriched terms, and assign those functions to all held out proteins in that cluster. We then compute the Jaccard similarity (Figure A12a,b,c) or F1 score using the top 10 terms (Figure A12d,e,f) between the assigned and true functional terms. The threshold of 10 is chosen based on the observation that most clusters have fewer than 10 enriched terms (Figure A12g,h,i). Across all three DREAM networks and diverse cluster sizes, ReCIPE improves upon the unconnected clusters in the function prediction task, yielding more functionally enriched clusters.

To test that the proteins ReCIPE is adding are meaningful and that the gains in performance are not just coming from adding more proteins, we compare ReCIPE with 50 random bootstraps adding a matched number of random proteins. As determined in the previous sections, we use settings of linear ratio = 10%, max proteins = 20. For each cluster, we show the average Jaccard similarity of 50 random bootstraps, compared to the similarity from the ReCIPE cluster (Figure A13, top). We also compute a statistical test of these results, using a Wilcoxon signed-rank test of the median position of the ReCIPE score vs. the random background, against a null hypothesis of 0.5 (i.e. ReCIPE clusters are no better or worse than the random distribution). Across all three networks, we find that ReCIPE significantly improves the Jaccard similarity of clusters (Figure A13, bottom). Before the ReCIPE step, an individual protein appears in at most one cluster. In A14 we show the distribution of cluster membership for each protein after ReCIPE; most proteins still appear in fewer than five clusters.

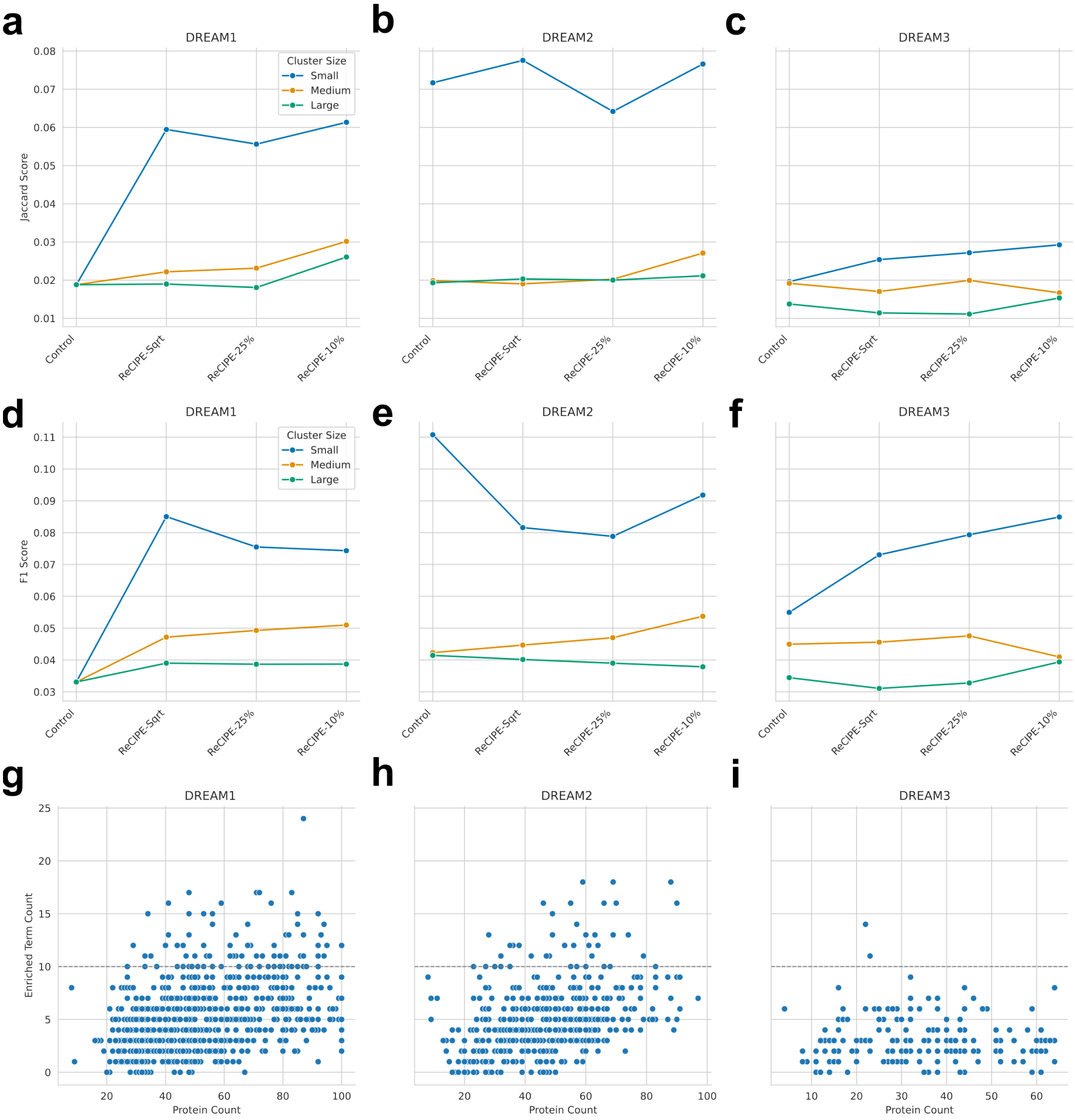


**Figure A12: ReCIPE improves percent enrichment of clusters on DREAM networks. (a,b,c)** ReCIPE improves Jaccard score of enriched terms on all three DREAM networks. Based on these experiments, we select a linear ratio of 10% to use for our analyses. **(d,e,f)** Using an alternative scoring, where we select the top 10 enriched terms and compute an F1 score, ReCIPE likewise improves on all three networks, except for small clusters in DREAM2. **(g,h,i)** Number of enriched terms for every cluster, plotted by number of proteins in that cluster. We use this to select the threshold of 10 enriched terms (dashed grey line) above, where 92.9%, 94.1% and 99.2% of clusters have fewer than or equal to 10 enriched terms.

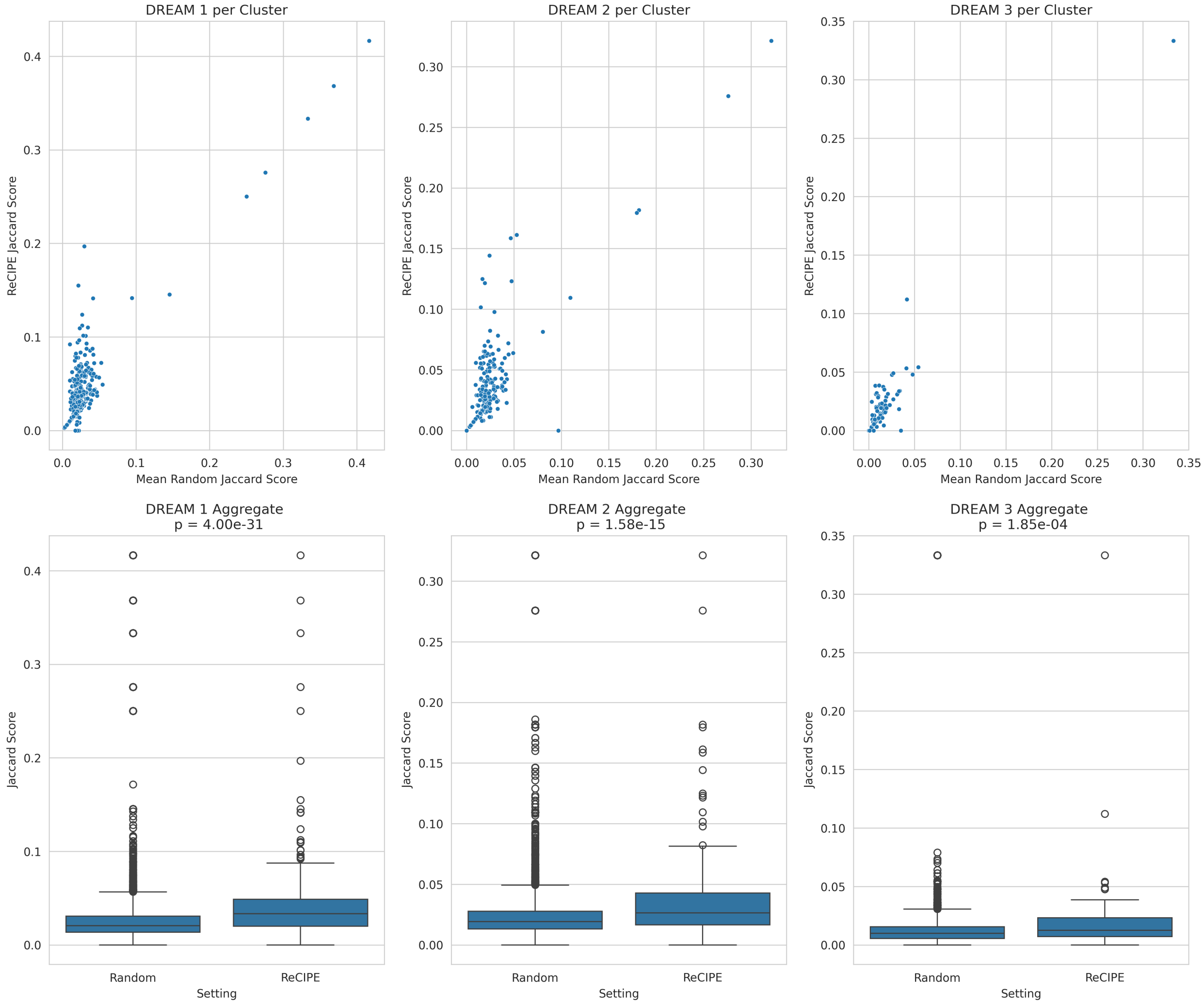


**Figure A13:** ReCIPE improvement in Jaccard similarity is not driven by larger cluster sizes.

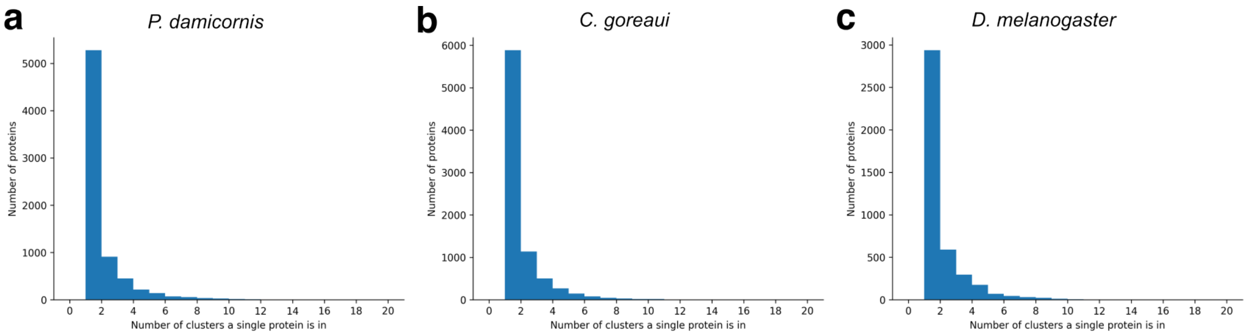


**Figure A14: Most proteins are still in a few clusters after ReCIPE.** Across all three species evaluated, ReCIPE adds a relatively small number of proteins back to clusters, maintaining them as largely non-overlapping. Most proteins appear in at most 5 clusters, with only a small handful of proteins appearing more than 8 times.

##### A.10.4 Ablation on inclusion order criteria

By default, ReCIPE prioritizes adding candidate nodes in order of increasing degree, to avoid adding non-specific hub nodes. Here, we consider four alternate methods for prioritizing node inclusion. These include:

- **Connected Components**, prioritizing first those candidates that connect to the most discrete components of the disconnected cluster
- **Incoming Edges**, prioritizing first those candidates that have the most edges to proteins in the disconnected cluster
- **Connected Components, Normalized**, the same as above but divided by the overall degree of the candidate node in the whole network
- **Incoming Edges, Normalized**, likewise the same, normalized by overall degree.

We evaluate each criteria in the same setting as in Section A.10.3, using the three DREAM networks and measuring the Jaccard Score and F1 score of cluster enrichment. We compute enrichment alternatively using either GSeaPy [111] or Func-E [110]. Scores are computed over all gene ontology terms, as well as each ontology (Biological Process, Molecular Function, Cellular Component). We thus have $3 \times 2 \times 4 = 24$ experimental settings; we drop the DREAM2 x Func-E x Molecular Function run due to a runtime error. We find that across the 23 settings analyzed, Jaccard and F1 score change minimally in response to inclusion criteria (Figure A15a,b,c), and that there is no significant difference in score. However, when we look at the relative ranks of each method in each experimental setting (Figure A15d,e,f), Incoming Edges, Normalized is frequently a top-ranked method, despite the small effect size. This motivates our change of the ReCIPE default to this new inclusion order.

#### A.11 Challenges and Limitations

Methods for remote homology detection [21] or structure-based search [112] are important steps in this direction, but network re-wiring between species [113] limits their applicability to genome-scale pathway analyses. Our study proves that the current generation of high-throughput PPI prediction methods are already accurate enough to enable network-wide functional genomics, but we stress that as these methods improve, so too will the accuracy and fidelity of downstream inference. There still remain several challenges and opportunities to improve the understanding of functional networks in non-model organisms. We note that our predicted networks are 6-10x as dense as the signal-to-noise ratio (SNR) estimated in true PPI networks (1:1000, [114]); thus we still likely have false positive edges and spurious connections. Although our downstream clustering approach will help with denoising, or a higher threshold

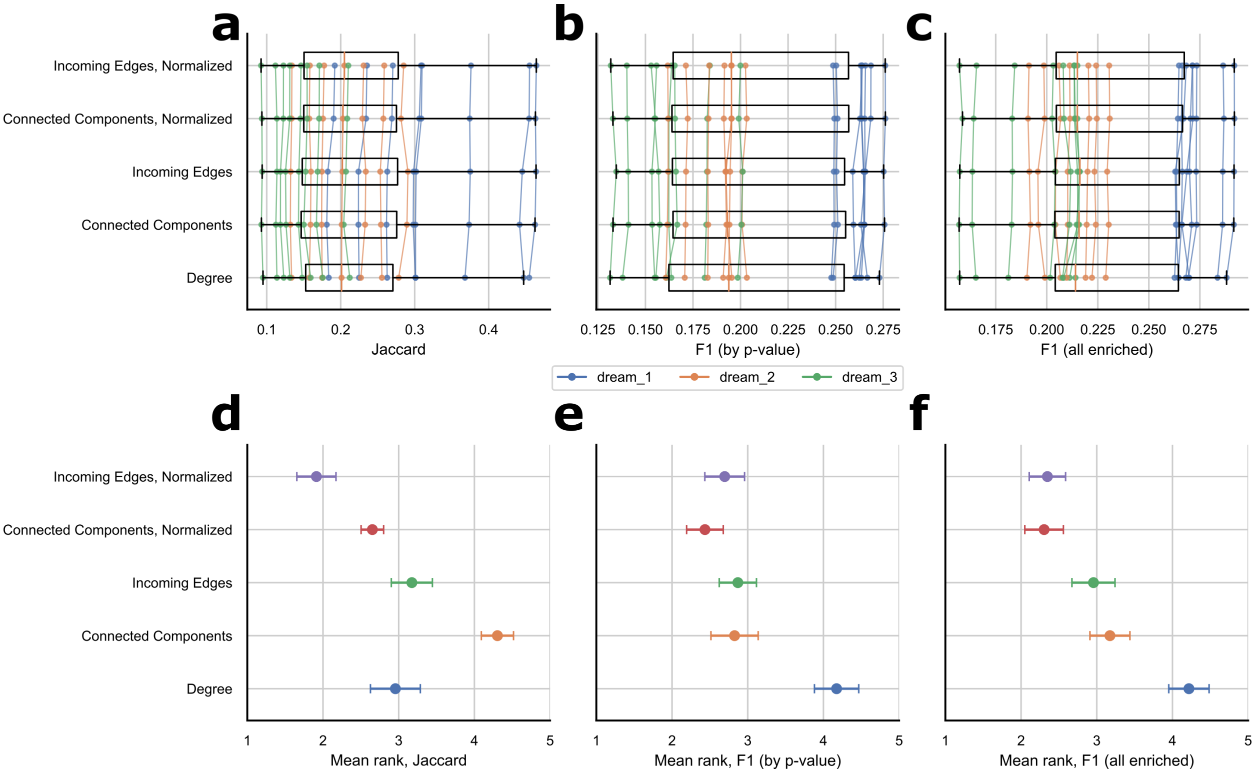


**Figure A15: Choice of inclusion priority criteria for ReCIPE.** By default, ReCIPE adds candidate nodes in order of increasing degree. We consider here four alternate ordering methods. **(a)** Functional enrichment scores are not significantly different depending on inclusion criteria across diverse experimental settings, suggesting that ReCIPE is robust to the order in which candidate proteins are added. **(b)** However, we do find that when we rank-order methods within each experimental setting, **Incoming Edges, Normalized** is consistently the highest performing method; we choose this as the new default for ReCIPE going forward.

for interaction could be selected to more closely match this SNR, improvements in network inference will ultimately have the largest impact on downstream performance. Structure-based PPI methods such as AlphaFold-Multimer [9] remain too slow for most labs to apply at interactome scale, but any sufficiently fast PPI prediction method can be substituted into PHILHARMONIC, and should improve its performance as the accuracy of these fast PPI prediction methods advance. Moreover, it is well known that protein interaction networks differ across tissues [64], and that sub-cellular localization likewise plays a role in protein interaction [115]. Our current approach assumes a single PPI network, and orthogonal information such as localization prediction [116], gene co-expression, or tissue type could help further refine analysis of functional networks.
